# Exosome tethering requires tetherin homodimerisation

**DOI:** 10.1101/2025.08.07.668892

**Authors:** Yağmur Yıldızhan, Adam M. Bourke, Hannah K. Jackson, Paul T. Manna, Roberta Palmulli, James R. Edgar

## Abstract

Exosomes are small extracellular vesicles that originate as intraluminal vesicles (ILVs) within multivesicular bodies (MVBs). Upon fusion of MVBs with the plasma membrane, ILVs are released into the extracellular environment as exosomes. While exosomes can diffuse away from their cells of origin, the expression of the antiviral restriction factor tetherin promotes their retention at the cell surface, thereby limiting their release into the extracellular milieu. Tetherin plays an analogous role in retaining other extracellular particles, including many enveloped viruses and midbody remnants. In this study, we examine the molecular basis of exosome tethering and define the mechanisms underlying tetherin-mediated retention. Using a combination of biochemical, live-cell and ultrastructural imaging approaches, we show that tetherin homodimerisation is essential for exosome tethering. Mutations in regions or motifs of tetherin predicted to impact tetherin traffic to ILVs have only minor impacts on exosome tethering, suggesting redundancy in the mechanisms of tetherin traffic to ILVs, and subsequently, to exosomes. Collectively, these findings provide the first molecular insights into the mechanism that govern exosome tethering.

## Introduction

Multivesicular bodies (MVBs) are specialized endosomal compartments that play a vital role in the sorting and trafficking of proteins, lipids, and other cellular components. A hallmark of MVBs is the presence of intraluminal vesicles (ILVs), which form through the inward budding of the MVB limiting membrane (Hirsch et al., 1968). This process encloses specific cargo, including proteins, RNA, and lipids, into ILVs, which are destined either for degradation within lysosomes (Felder et al., 1990) or for release into the extracellular environment (Harding et al., 1983; Pan and Johnstone, 1983; Raposo et al., 1996). ILVs can be generated through both ESCRT-dependent (Hurley and Hanson, 2010; Raiborg and Stenmark, 2009) and ESCRT-independent pathways (Edgar et al., 2014; Stuffers et al., 2009; Theos et al., 2006; van Niel et al., 2011), and different subtypes of ILVs coexist within single MVBs (Edgar et al., 2014). Once MVBs are formed, they follow one of two primary routes: fusion with the lysosome for degradation of ILVs, or fusion with the plasma membrane leading to the release of ILVs into the extracellular space, where they are designated as exosomes (Raposo et al., 1996).

Exosomes play a crucial role in intercellular communication by transferring bioactive molecules between cells, influencing physiological and pathological processes such as immune modulation (Bobrie et al., 2011; Muntasell et al., 2007; Nolte-’t Hoen et al., 2009; Théry et al., 2002), skin pigmentation (Lo Cicero et al., 2015; Prospéri et al., 2024), tumour progression (Hoshino et al., 2013; Palmulli et al., 2025; Peinado et al., 2012), and the propagation of pathogenic proteins including Prions (Fevrier et al., 2004), beta amyloid (Rajendran et al., 2006) and alpha synuclein (Emmanouilidou et al., 2010).

Historically, exosomes were believed to be released from cells to diffuse freely into the extracellular space to act on cells in *trans*. Our previous work, however, demonstrated instead that exosomes can also be retained at the plasma membrane of the producer cell through a process mediated by the antiviral restriction factor tetherin (*Bst2*/CD317) (Edgar et al., 2016). The observation that exosomes remain tethered to the plasma membrane highlighted additional possible functions for exosomes, and largely that tethered exosomes could provide a high local concentration of molecules at the cell surface that are not subject to lateral diffusion in the same way as those at the plasma membrane. Tetherin is widely expressed, and its expression is highly sensitive to interferon (Neil et al., 2007). Although numerous published micrographs show clusters of exosomes at the surface of cells, the function of tethered exosomes remain largely unknown. Tethered exosomes have recently been found to contribute to the degradation of the pericellular extracellular matrix in breast cancers (Palmulli et al., 2025), and this finding may explain the previously unresolved link between tetherin overexpression in metastatic cancers and metastasis formation (Mahauad-Fernandez and Okeoma, 2017; Mahauad-Fernandez et al., 2014).

In addition to exosomes, tetherin mediates the retention of other extracellular particles, including enveloped viruses (Kaletsky et al., 2009; Lopez et al., 2010; Mansouri et al., 2009; Neil et al., 2008; Stewart et al., 2023; Wang et al., 2014) and midbody remnants (Presle et al., 2021), highlighting its broad role in regulating the retention and release of small membranous particles that are generated by outward budding. The formation of exosomes, enveloped viruses, and midbody remnants occurs via membrane budding events directed away from the cytosol - either toward the extracellular space or into the lumen of endosomes. These outward budding processes utilize specialized molecular machinery. Cytokinesis, which results in the formation of midbody remnants, as well as the biogenesis of a subset of ILV/exosomes and enveloped viruses, involves the Endosomal Sorting Complex Required for Transport (ESCRT) machinery (McDonald and Martin-Serrano, 2009). Whether tetherin also restricts other forms of extracellular vesicles, such as ectosomes and apoptotic bodies, remains unclear.

Human tetherin consists of a short N-terminal cytosolic domain, a transmembrane domain, and an extracellular coiled-coil domain, which is anchored to the membrane via a C-terminal GPI anchor (Kupzig et al., 2003). The extracellular coiled-coil domain contains three conserved cysteine residues essential for homodimer formation, a key requirement for the retention of both Human Immunodeficiency Virus-1 (HIV-1) particles (Perez-Caballero et al., 2009) and midbody remnants (Presle et al., 2021). While the structural features of tetherin are well characterized in the context of viral restriction, their role in exosome tethering remains poorly understood. Determining which structural elements are required for exosome retention is crucial for elucidating the molecular mechanisms that regulate exosome release and surface accumulation.

In this study, we aim to elucidate the molecular mechanism underlying tetherin-mediated exosome tethering. Using a combination of molecular, biochemical, live-cell and ultrastructural imaging approaches we demonstrated that tetherin homodimer formation is crucial for exosome tethering. By generating tetherin mutants that disrupt specific functions, such as dimerisation, ubiquitination, and the addition of a GPI anchor, we assessed their impact on exosome retention. We also examined the molecular machinery that traffics tetherin to ILVs, and which features of tetherin mediate this trafficking. Together, our results provided the first mechanistic insights into how tetherin controls exosome retention at the plasma membrane and clarify the structural determinants underlying this function.

## Materials and methods

### Cell lines and standard culture conditions

WT HeLa cells were a gift from Professor Scottie Robinson (CIMR, University of Cambridge, UK). Bst2KO HeLa cells were previously described (Edgar et al., 2016). HEK293T cells were purchased from ATCC. MelJuSo cells were a gift from Dr Claire Shannon-Lowe (University of Birmingham). All cell lines were cultured in DMEM supplemented with 10% fetal bovine serum and 1% Penicillin/Streptomycin, and maintained at 37°C in a 5% CO_2_ incubator Bst2KO MelJuSo cells were generated using px330-Bst2 and the gRNA (5’-GCTCCTGATCATCGTGATTC**TGG)** as previously described (Edgar et al., 2016). Bst2KO MelJuSo cells were bulk sorted by FACS, expanded, and verified by western blot, flow cytometry and immunofluorescence (as shown in Figure 5).

All cells were regularly screened for mycoplasma using a PCR Mycoplasma detection kit (Abcam, ab289834).

### Cloning

cDNAs for Bst2-WT-HA, Bst2-C3A-HA, Bst2-K18A-HA, Bst2-Nterm-HA, Bst2-GPI-HA were a gift from Professor George Banting (University of Bristol, UK) and are previously described (Billcliff et al., 2013). HA-tagged tetherin cDNAs were PCR’ed to add NotI and XhoI restriction sites and cloned into the retroviral vector, pQCXIH.

cDNAs for Bst2-WT-mScar and Bst2-C3A-mScar were synthesised by GeneWiz (Azenta Life Sciences) and cloned to pQCXIH between AgeI and BamHI restriction sites. mScarlet cDNA was cloned between position Y154/P155 of human *Bst2* and separated by short, flexible (Gly/Ser) linkers.

pCMV-Sport6-CD63-pHluorin was a gift from Dr Michiel Pegtel (Amsterdam UMC, Netherlands).

All plasmids were confirmed by Sanger sequencing. Sequences are provided in Table 1.

**TABLE 1.**
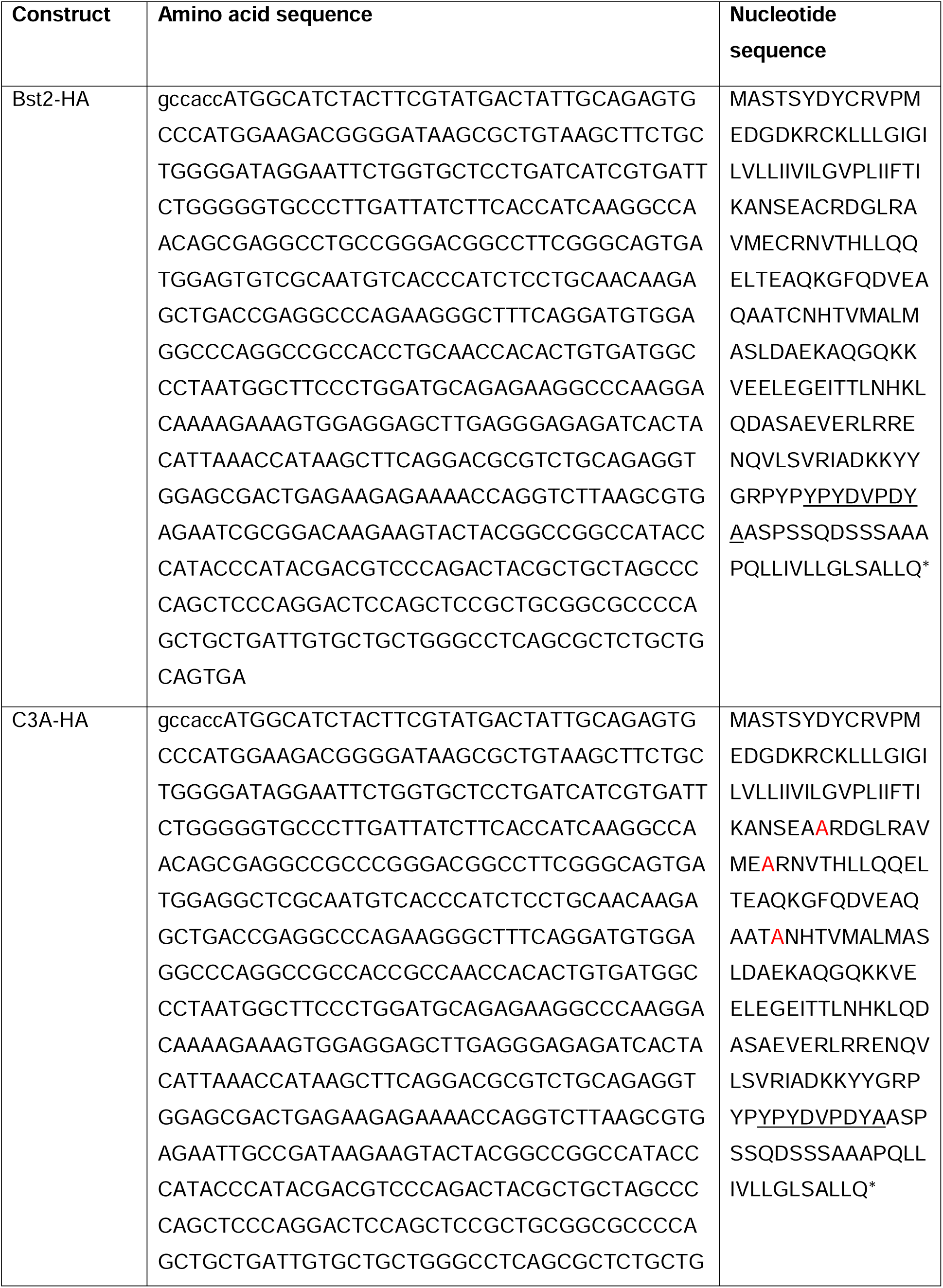
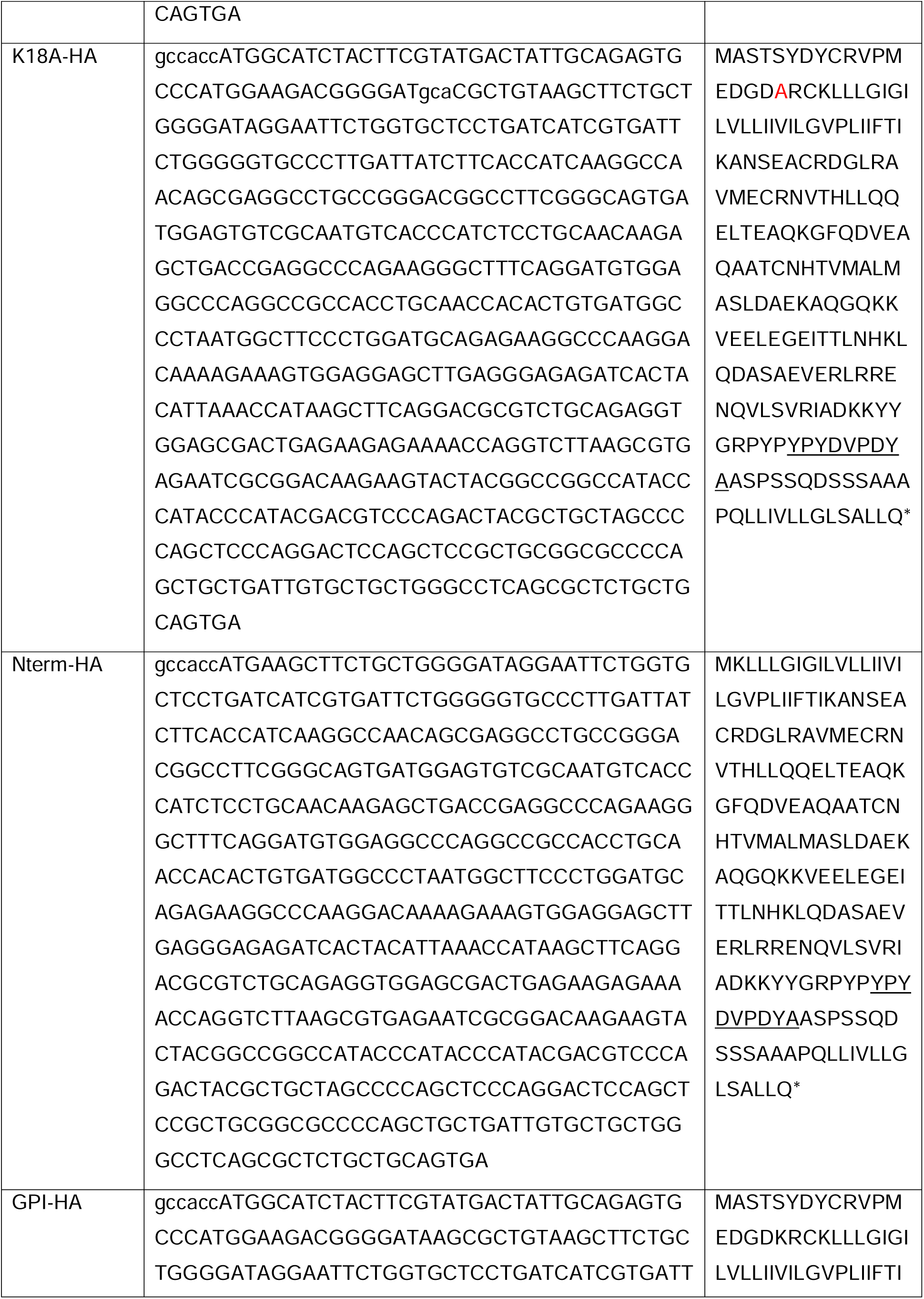
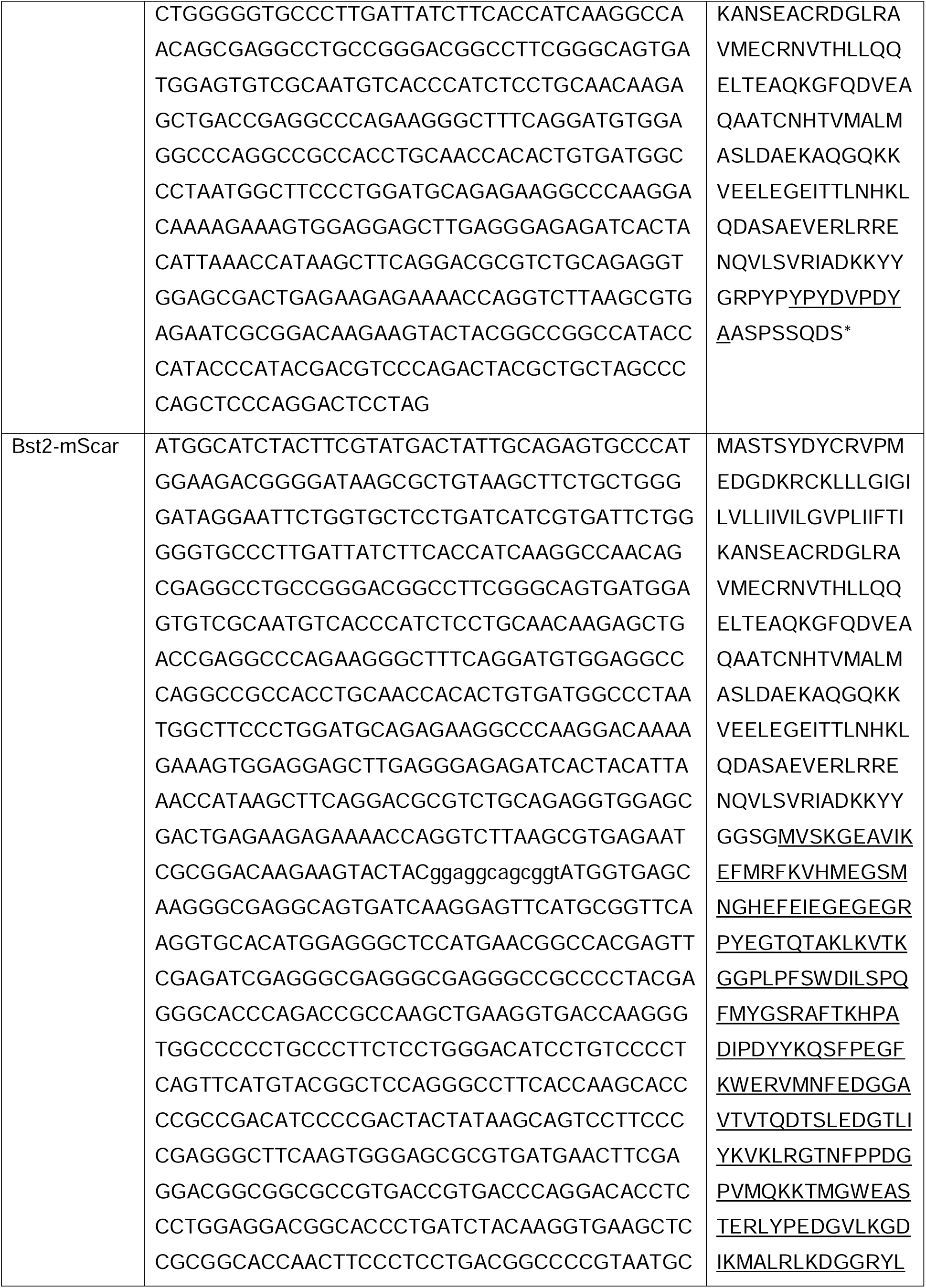
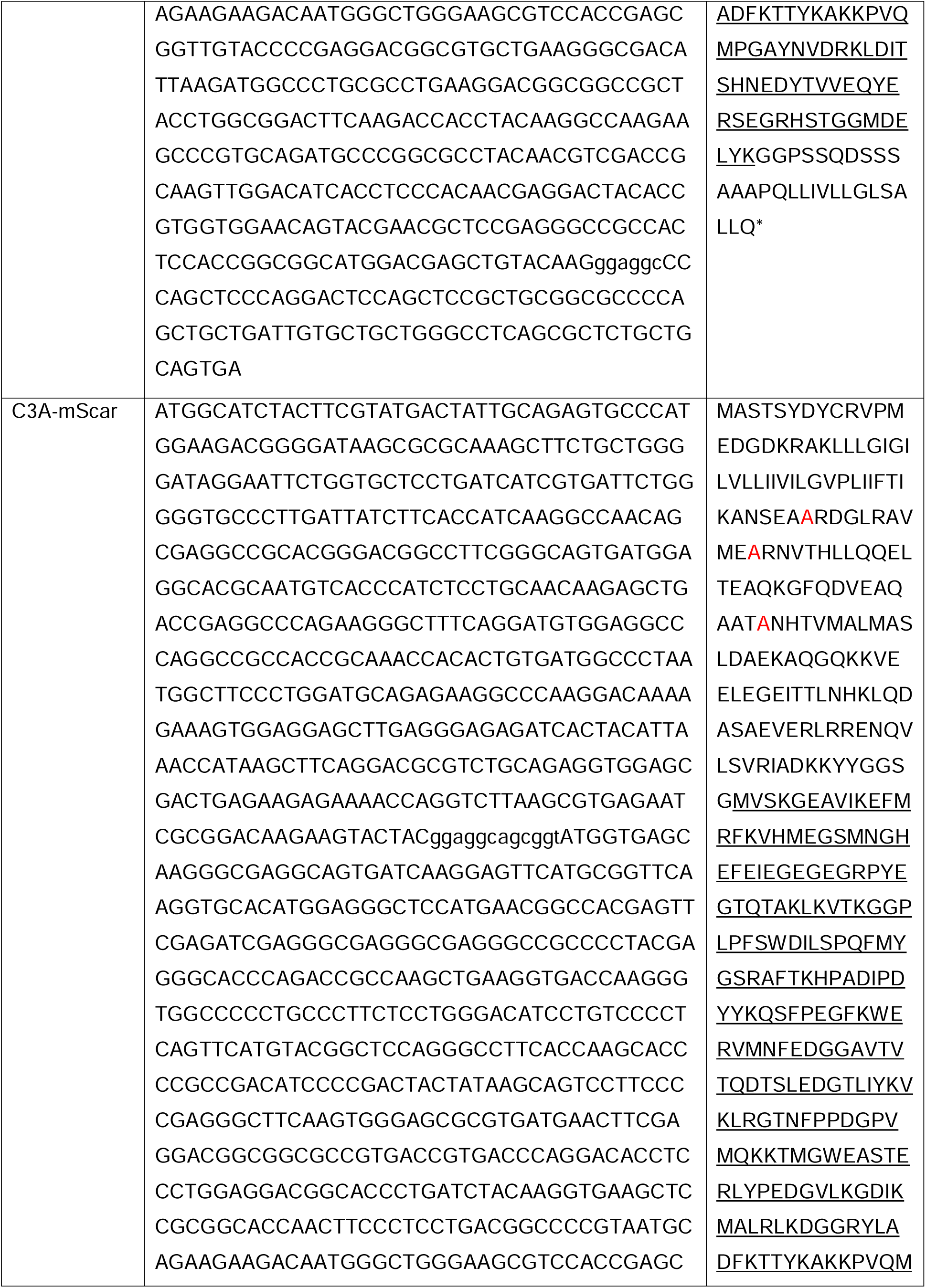
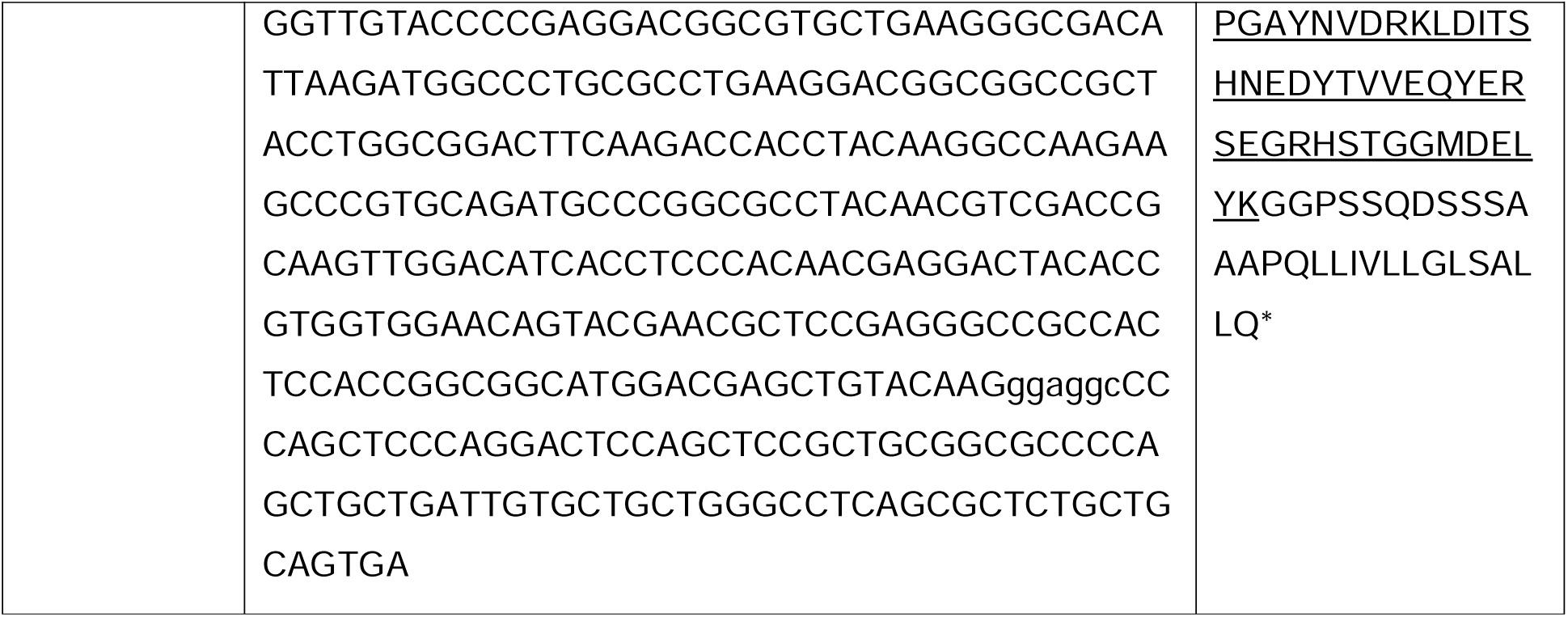

### Generation of tetherin rescue cell lines

Retroviral pQCXIH vectors were co-transfected into HEK293T cells with pMD.GagPol and pMD.VSVG packaging plasmids using TransIT-293 (Mirus Bio, USA). Viral supernatants were collected 48 hours after transfection, passed through 0.45 μm filters to remove dead cells and debris. Recipient cells were transduced by ‘spinfection’-viral supernatants were centrifuged at 1800 rpm in a benchtop centrifuge at 37°C for 70 mins to enhance viral transduction. Four days post-transduction, cells were subject to antibiotic selection with 200 µg/mL Hygromycin B.

HA- and mScar-tetherin stable cell lines were generated by retroviral transduction of Bst2KO HeLa or Bst2KO MelJuSo cells.

### Antibodies

The following primary antibodies were used: anti-tetherin (rabbit monoclonal, Abcam, ab243230, WB 1:2500, IF 1:200, immunoEM 1:20), anti-EEA1 (mouse monoclonal, BD Transduction, 610457, IF 1:200), anti-VPS35 (mouse monoclonal, Santa Cruz, B-5, IF 1:200), anti-CD63 (mouse monoclonal, BioLegend, H5C6, WB 1:1000, IF 1:200), anti-LAMP1 (mouse monoclonal, BioLegend, H4A3, IF 1:200), anti-CIMPR (mouse monoclonal, Abcam, ab2733, IF 1:200), anti-GM130 (mouse monoclonal, BD Biosystems, 35/GM130, IF 1:200, WB 1:1000), anti-GAPDH (mouse monoclonal, Proteintech, 1E6D9, WB 1:10,000), anti-LBPA (mouse monoclonal, Millipore, 6C4, IF 1:200), anti-HA (rat monoclonal, Roche, 3F10, IF 1:200), anti-HA (rabbit monoclonal, Cell Signalling, C29F4, WB 1:1000, immunoEM 1:30), anti-ALIX (mouse monoclonal, GeneTex, 3A9, WB 1:1000), anti-Syntenin (rabbit monoclonal, Abcam, ab133267, WB 1:1000), anti-Tsg101 (mouse monoclonal, GeneTex, 4A10, WB 1:1000), anti-CD9 (rabbit monoclonal, Abcam, ab92726, WB 1:500), anti-Hrs (rabbit polyclonal, a gift from Prof Sylvié), anti-HLA-DR (rabbit monoclonal, abcam, ab259258, WB 1:1000). PE-conjugated anti-Bst2 (mouse monocloncal, BioLegend, RS38E, FC 1:200), APC-conjugated anti-HA (Mouse monocloncal, Miltenyi, CG8-1F3.3.1, FC 1:200), Secondary antibodies used in the study were: anti-rat IgG Alexa-488 (goat, Molecular Probe, A11006, IF 1:200), anti-rabbit IgG Alexa 488/555 (goat, ThermoFisher, IF 1:200) and anti-mouse IgG Alexa-488/555 (goat, ThermoFisher, IF 1:200). For Western blotting, anti-mouse IRDye 800CW (goat, LI-COR, 926-32210, WB 1:10,000) and anti-rabbit IRDye 680RD (goat, LI-COR, 926-68071, WB 1:10,000) were used.

### Reagents and small molecules

Leupeptin Hemisulphate was purchased from Cambridge Bioscience (HY-18234A-1ml), recombinant human IFN-gamma from PeproTech (300-02-100UG), Lipofectamine 2000 from Invitrogen (11668-030), and Bafilomycin A1 was from Sigma-Aldrich (B1793).

### Western blotting

Whole cells or exosome-enriched pellets were lysed in lysis buffer (1% Triton-X100, 1mM EDTA, 150mM NaCl, 20 mM Tris pH 7.5) supplemented with 1 x cOmplete EDTA-free Protease Inhibitor Cocktail (11836170001, Roche) on ice for 30 min with regular vortexing. Lysates were centrifuged at 14,000 ×*g* for 15 min at 4°C. Resultant lysates were mixed with 4x NuPage LDS sample buffer (ThermoFisher), boiled at 65°C or 95°C for 10 min and subjected to SDS-PAGE on NuPage 4-12% Bis-Tris precast gels (ThermoFisher). Samples were transferred to PVDF membranes. Membranes were blocked in 5% non-fat dried milk in PBS with 0.1% Tween-20 (PBS-T) for 30 minutes at room temperature. Then, they were incubated with primary antibodies overnight at 4°C, followed by secondary antibodies for 1 hour at room temperature. Finally, the membranes were imaged with an Odyssey CLx system (LI-COR).

Signal intensities were quantified using ImageJ Fiji software, with quantifications based on at least three independent experiments. The presented immunoblots are representative of these experiments.

### Immunofluorescence microscopy

Cells were seeded to 13mm glass coverslips and left overnight to adhere. The following day, cells were fixed with 4% PFA/PBS. Cells were quenched with 15 mM glycine/PBS and permeabilised using 0.1% saponin/PBS. Blocking and subsequent antibody labelling were performed with 1% BSA supplemented with 0.01% saponin/PBS. Nuclei were labelled with Hoechst. Cells were mounted to slides with mounting medium and images were acquired using a LSM700 confocal microscope (63×, 1.4nA, oil immersion objectives; ZEISS).

### Colocalisation analysis

ImageJ Fiji version 2.3.0 was used for colocalisation analysis. Mander’s correlation coefficient between two channels was quantified using JACoP plugin of ImageJ Fiji software. At least twenty cells from three independent experiments were analyzed for each condition.

### Immunofluorescence analysis – CD63 particle density

Immunofluorescence images were analysed using ImageJ Fiji version 2.3.0 to quantify CD63-positive particle count over cell area. Images were converted to 8-bit and background-subtracted. A consistent threshold was applied across all samples to segment CD63-positive puncta. The “Analyse Particles” function was used to count individual puncta. For each image, the cell area was manually outlined, and CD63 particle counts were normalized to the cell area. Particle density was calculated as: (number of CD63-positive particles / cell area in µm²) × 100. This yielded density values expressed as particles per 100 µm. Data were analysed using GraphPad Prism. Group comparisons were performed using ordinary one-way ANOVA followed by Tukey’s multiple comparisons test. *p* > 0.05 (ns), *p* ≤ 0.05 (*), *p* ≤ 0.01 (**), *p* ≤ 0.001 (***) and *p* ≤ 0.0001 (****).

### Flow cytometry

Cells were gently trypsinised and centrifuged at 350 ×*g* for 4 min at 4°C. Cell pellets were resuspended in FACS buffer (PBS containing 1% BSA and 1 mM EDTA), transferred to 96-well plates, and washed once. For intracellular staining, cells were fixed, permeabilised with 0.1% saponin in FACS buffer, then incubated with fluorophore-conjugated antibodies diluted in 0.05% saponin/FACS buffer for 45 min on ice. For surface staining, cells were incubated with fluorophore-conjugated antibodies in FACS buffer for 45 min on ice. For staining with unconjugated primary antibodies, cells were incubated with the primary antibody for 45 min, washed, and subsequently incubated with Alexa-conjugated secondary antibodies. Prior to data acquisition, cells were washed twice with cold PBS. Fluorescence intensity was analysed using a Cytoflex LX flow cytometer (Beckman Coulter) equipped with six lasers (488, 561, and 638 nm lasers were used).

### siRNA-mediated knockdown of Hrs

Cells were transfected with ON-TARGET*plus* siHrs pool (*Hgs*) (Horizon Discovery) using Lipofectamine RNAiMax (ThermoFisher) on days 1 and 2 and harvested on day 4. Non-targeting siRNA were used as a control (ON-TARGET*plus* Non-targeting Pool, Horizon Discovery).

### Exosome-enriched preparations

Exosome-enriched preparations were generated from cell culture supernatants by differential ultracentrifugation following the protocol described in (Théry et al., 2006). Briefly, cells were cultured in equal numbers on three T175 tissue culture flask containing 30 mL of media. To induce exosome release, cells were treated with 100 nM Bafilomycin A1 for 16 hours. The culture supernatants were collected and centrifuged at 300 ×*g* for 10 minutes to remove large cell debris. The supernatants were then centrifuged at 2000 ×*g* for 10 minutes to eliminate smaller cell debris. Subsequently, the cleared supernatants were ultracentrifuged at 10,000 ×*g* for 30 minutes using a Type 70 Ti Rotor (Beckman Coulter) to remove larger extracellular vesicles. The resulting supernatants were collected and ultracentrifuged again at 100,000 ×*g* for 70 minutes to pellet exosomes. These pellets were washed in PBS and re-pelleted by ultracentrifugation at 100,000 ×*g* for an additional 70 minutes. The final exosome-enriched pellets were either used immediately or stored at -80°C for future experiments.

### Conventional transmission electron microscopy

Cells were grown on Thermanox coverslips (Nunc) and left overnight to adhere. Cells were fixed with 4% PFA/5% glutaraldehyde/0.1 M cacodylate buffer (pH 7.4) at a 1:1 ratio with culture medium (final concentration 2% PFA/2.5% glutaraldehyde) for 30 minutes. Cells were washed with 0.1 M cacodylate before being post-fixed with 1% osmium tetroxide: 1.5% potassium ferrocyanide. Samples were contrast enhanced using 1% tannic acid/0.1M cacodylate buffer. Coverslips were washed with water before being dehydrated with an ethanol series. Epoxy propane was used to infiltrate cells (CY212 Epoxy resin (Agar Scientific):propylene oxide (Agar Scientific) before being infiltrated with neat CY212 Epoxy resin. Coverslips were mounted to pre-baked resin stubs and polymerised overnight at 65°C. Thermanox coverslips were removed with a heat block. 70 nm sections were cut using a Diatome diamond knife mounted to an ultramicrotome and were collected to formvar coated grids before being stained with lead citrate. An FEI Tecnai transmission electron microscope was used to visualise samples at an operating voltage of 80kV.

### Tethered exosome quantification

The entire plasma membrane of cells was screened and the number of extracellular, membranous vesicles with a diameter <150nm were counted. The length of the corresponding plasma membrane was calculated, and the resultant number of exosomes per µm plasma membrane generated. Exosome frequency analysis was excluded from areas where cells were within 500nm of another cell, to minimise any potential impact from adjacent cells physically blocking exosome release.

To exclude any bias from the height of the ultrathin section, at least two different sections (of different relative height) per biological replicate were analysed, and two independent biological repeats were performed. At least 5 independent cells per section were quantified (at least 20 cells analysed per condition total).

### Cryo Immunogold transmission electron microscopy

Cells were fixed with 2% PFA/0.1M phosphate buffer overnight at 4°C. Cells were scraped in 12% gelatine and immediately pelleted by centrifugation. Cell pellets were infiltrated by PVP/Sucrose before being mounted to aluminium pins, frozen in liquid nitrogen and cut at - 90°C using a Diatome Diamond knife attached to a cryo ultramicrotome.

Thawed cryosections were labelled with anti-HA or anti-tetherin antibodies, prior to incubation with 10nm Protein-A gold. Labelled sections were stained using uranyl acetate:methyl cellulose prior to imaging using an FEI Tecnai transmission electron microscope at an operating voltage of 80kV.

### Nanoparticle Tracking Analysis (NTA) with Zetaview

Exosome-enriched samples were diluted in PBS to 1=mL and pre-tested to ensure optimal particle concentrations (140–200 particles/frame). Measurements were performed using the ZetaView system (Particle Metrix). Each sample was analysed in three cycles, scanning 11 positions and recording 60 frames per position (video setting: high). Acquisition settings included autofocus, camera sensitivity 92.0, shutter 70, scattering intensity 4.0, at 25=°C.

Videos were analysed using ZetaView Software (v8.02.31) with analysis parameters: particle size range 5-1000 and minimum brightness 20. All samples exceeded 1000 completed tracks to ensure robust quantification.

### Nanoparticle Characterisation

For DAISY analysis (HoltraAB, Gothenburg, Sweden), samples were diluted to approximately 5×10^8^ particles per mL in PBS. Exosome-enriched samples were applied to microfluidic chips (Topaz, design 180, Topaz polymer, ChipShop, Jena, Germany). From each recorded particle, averaged optical properties were extracted in both digital holographic and interferometric scattering (iSCAT) imaging modes simultaneously.

### TIRF microscopy

Total Internal Reflection Fluorescence (TIRF) microscopy was used to monitor exosome fusion events in MelJuSo cells stably expressing either Bst2-mScarlet or C3A-mScarlet. Cells were seeded on 35=mm MatTek glass-bottom dishes (Part No.: P35G-1.5-10-C, MatTek, USA) and transiently transfected with CD63-pHluorin 24 hours prior to imaging. Live-cell imaging was conducted using a ZEISS Elyra 7 microscope equipped with an alpha Plan-Apochromat 63×/1.46 Oil Korr M27 TIRF objective. Imaging was performed in phenol red-free complete culture medium maintained at 37=°C and 5% CO_₂_. Time-lapse sequences were acquired at 2 or 3=Hz for a total duration of 1 minute.

Fusion events -interpreted as exosome fusion or retention at the plasma membrane - were identified manually as sudden, localised increases in fluorescence intensity. Fusion activity was quantified as the number of events per μm² of cell surface. The duration of each event was defined as the time elapsed from the peak fluorescence intensity until the signal returned to baseline. 20 cells were analysed across at least three biologically independent experiments to ensure reproducibility.

## Results

### Tetherin distributes throughout the endolysosomal pathway

Tetherin localises to a number of organelles, enabling it to restrict various extracellular particles, including enveloped viruses, midbody remnants and exosomes. The localisation of tetherin to these distinct organelles allows its incorporation into the membrane of the subsequent extracellular particle. Tetherin restricts different classes of enveloped viruses that bud from distinct organelles - for example, retroviruses (such as HIV-1) bud predominately from the plasma membrane (Morita and Sundquist, 2004; Welsch et al., 2007), and Coronaviruses (such as SARS-CoV-2) bud predominately into ERGIC/Golgi organelles (Goldsmith et al., 2004; Klein et al., 2020; Salanueva et al., 1999). Tetherin distribution to both the plasma membrane and to biosynthetic organelles allows it to restrict both classes of enveloped viruses. Similarly, tetherin localisation at the plasma membrane allows it to restrict midbody remnants (Presle et al., 2021). In order for tetherin to tether exosomes, it is first trafficked to ILVs of MVBs (Edgar et al., 2016) – the precursors of exosomes. The trafficking of tetherin between these organelles is therefore key to promoting its incorporation into enveloped viruses, midbody remnants and exosomes, and vital for its ability to tether such particles.

To understand the distribution of tetherin within the endolysosomal pathway we performed confocal microscopy. Endogenous tetherin displays colocalisation with early endosomes (EEA1), late endosomes (VPS35 / CD63), and lysosomes (LAMP1) (**Figure 1A**). Tetherin showed highest colocalisation with VPS35 (**Figure 1B**) – a component of the retromer complex which traffics cargos in a retrograde direction from endosomes towards the biosynthetic pathway (Seaman et al., 1998).

**Figure 1.**
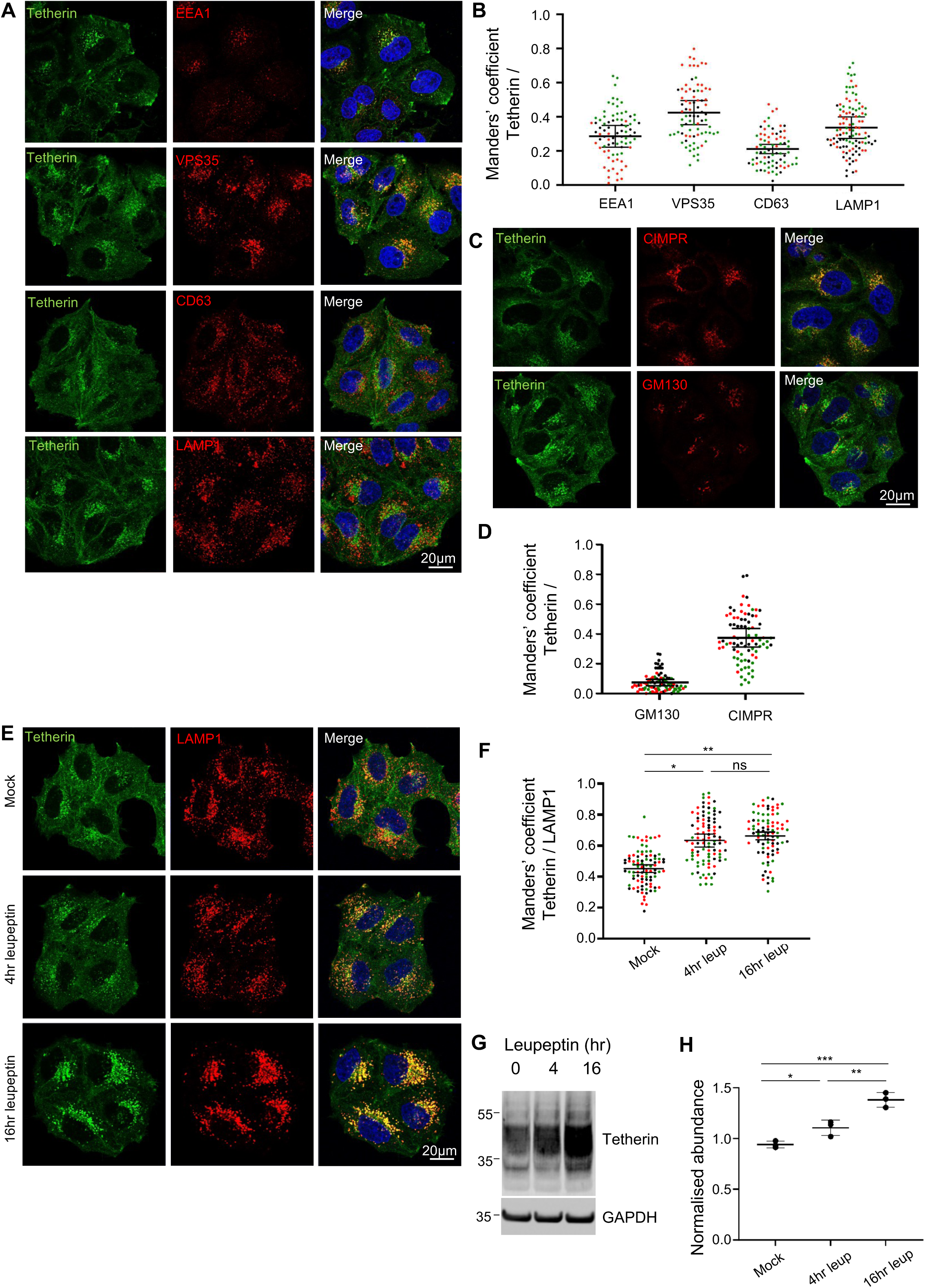
**A.** Representative confocal images showing colocalisation of tetherin with endosomal markers in WT HeLa cells, stained with DAPI (blue), anti-tetherin (red), and one of the following endosomal markers (green): EEA1 (early endosomes), VPS35 (retromer complex), CD63 (late endosomes/MVBs, or LAMP1 (lysosomes). Scale bars, 20=μm. **B.** Quantification of colocalisation between tetherin and endosomal markers. Manders’ correlation coefficients between tetherin and the endosomal markers EEA1, VPS35, CD63, LBPA, and LAMP1 in WT HeLa cells. Data represent three independent biological replicates (*n* > 74 cells), each dot corresponds to an individual cell; red, green, and black colours represent separate biological replicates. Bars indicate mean ± SEM. Mean values: tetherin/EEA1 = 0.28 ± 0.06; tetherin/VPS35 = 0.42 ± 0.07; tetherin/CD63 = 0.21 ± 0.03; tetherin/LBPA = 0.18 ± 0.03; tetherin/LAMP1 = 0.34 ± 0.06. **C.** Representative confocal images showing colocalisation of tetherin with biosynthetic pathway markers in WT HeLa cells, stained with DAPI (blue), anti-tetherin (red), and either GM130 (Golgi apparatus) or CIMPR (trans-Golgi network/early endosomes) (green). Scale bars = 20=μm. **D.** Quantification of colocalisation between tetherin and biosynthetic pathway markers. Manders’ correlation coefficients between tetherin and GM130 or CIMPR in WT HeLa cells. Data represent three independent biological replicates (*n* > 82 cells), each dot corresponds to an individual cell; red, green, and black colours represent separate biological replicates. Bars indicate mean ± SEM. Mean values: tetherin/GM130 = 0.08 ± 0.02; tetherin/CIMPR = 0.38 ± 0.06. **E.** Representative confocal images showing colocalisation of tetherin and LAMP1 following leupeptin treatment in WT HeLa cells. Cells were incubated with 50=μg/mL leupeptin for 4=hr or 16=hr, fixed, and stained with DAPI (blue), anti-tetherin (green), and anti-LAMP1 (red) antibodies. Scale bars = 20=μm. **F.** Quantification of colocalisation between tetherin and LAMP1 in WT HeLa cells following leupeptin treatment. Manders’ correlation coefficients between tetherin and LAMP1 are shown. Data represent three independent biological replicates (*n* > 87 cells per condition), each dot represents an individual cell; red, green, and black dots denote separate biological replicates. Bars indicate mean ± SEM. Mean values: mock = 0.45 ± 0.03; 4=hr = 0.64 ± 0.04; 16=hr = 0.66 ± 0.03. Statistical analysis was performed using ordinary one-way ANOVA followed by Tukey’s multiple comparisons test. Mock vs 4=hr (*p* = 0.0161), mock vs 16=hr (*p* = 0.0081), 4=hr vs 16=hr (*p* = 0.8046). *p* > 0.05 (ns), *p* ≤ 0.05 (*), *p* ≤ 0.01 (**), *p* ≤ 0.001(***) and *p* ≤ 0.0001 (****). **G.** Representative western blot of tetherin expression in WT HeLa cells following leupeptin treatment. WT HeLa cells were either untreated (mock) or incubated with 50=μg/mL leupeptin for 4 or 16 hr. Total cell lysates were analysed by western blotting using anti-tetherin antibodies in non-reduced conditions. GAPDH was used as a loading control. **H.** Quantification of tetherin protein levels in mock WT HeLa cells, or cells treated with leupeptin (4 hr or 16 hr). Tetherin abundance was assessed by western blotting and normalised to GAPDH. Each dot represents the normalised tetherin level from one independent experiment (*n* = 3). Bars indicate mean ± SEM (mock = 1.03=± 0.02, 4 hr = 1.21±=0,05, 16 hr = 1.52=±=0.05). Statistical analysis was performed using one-way ANOVA with Tukey’s multiple comparisons test. *p* > 0.05 (ns), *p* ≤ 0.05 (*), *p* ≤ 0.01 (**), *p* ≤ 0.001(***) and *p* ≤ 0.0001 (****). Mock vs 4hr leupeptin (*p* = 0.0445), mock vs 16hr leupeptin (*p* = 0.0004).

Given that cholesterol-rich MVBs are more prone to fuse with the plasma membrane than lysosomes (Möbius et al., 2003), and that tetherin contains a C-terminal GPI anchor which its retention in cholesterol-rich domains (Billcliff et al., 2013; Brown and Rose, 1992), we examined whether tetherin were positive for cholesterol. Many tetherin puncta colocalised with cholesterol (**Supplementary** Figure 1A), but interestingly, tetherin localised away from the LBPA, a phospholipid specifically enriched in late endosomes and lysosomes (**Supplementary** Figure 1B).

Upon reaching endosomes, a proportion of tetherin undergoes retrograde traffic and this enables tetherin levels to be restored in the biosynthetic pathway and plasma membrane (Stewart et al., 2023). In addition to localising to punctate organelles, tetherin displays some perinuclear staining which strongly colocalises with CIMPR, but not with the cis-Golgi marker GM130 (**Figure 1C, D**).

Late endosomes can either fuse with terminal lysosomes to form endolysosomes (Bright et al., 1997), or with the plasma membrane to release ILVs to the extracellular environment where they become termed exosomes (Raposo et al., 1996). To determine whether tetherin-positive endosomes are selectively fated for fusion with the plasma membrane (and therefore escape from lysosomal degradation) we examined the effect of Leupeptin, a lysosomal protease inhibitor, on tetherin trafficking and abundance. Immunofluorescence microscopy revealed that treatment with Leupeptin led to an increased abundance of tetherin in LAMP1-positive organelles (**Figure 1E, F**), and Western blotting confirmed elevated total tetherin (**Figure 1G, H**). These findings collectively indicate that tetherin traffics throughout the endolysosomal pathway and that tetherin-positive late endosomes are not fated specifically for fusion with the plasma membrane.

### Tetherin homodimer formation is required for exosome tethering

While the molecular mechanisms of tetherin-mediated virus and midbody remnant restriction have been elucidated, the mechanisms governing exosome tethering remain undefined (**Figure 2A**). One possibility is that individual tetherin molecules span opposing membranes – branching between an exosome and the plasma membrane (*branch model*). Alternatively, individual tetherin molecules may be embedded in a single membrane but form disulfide-linked homodimers with tetherin molecules in an opposing membrane (*homodimer model*).

**Figure 2.**
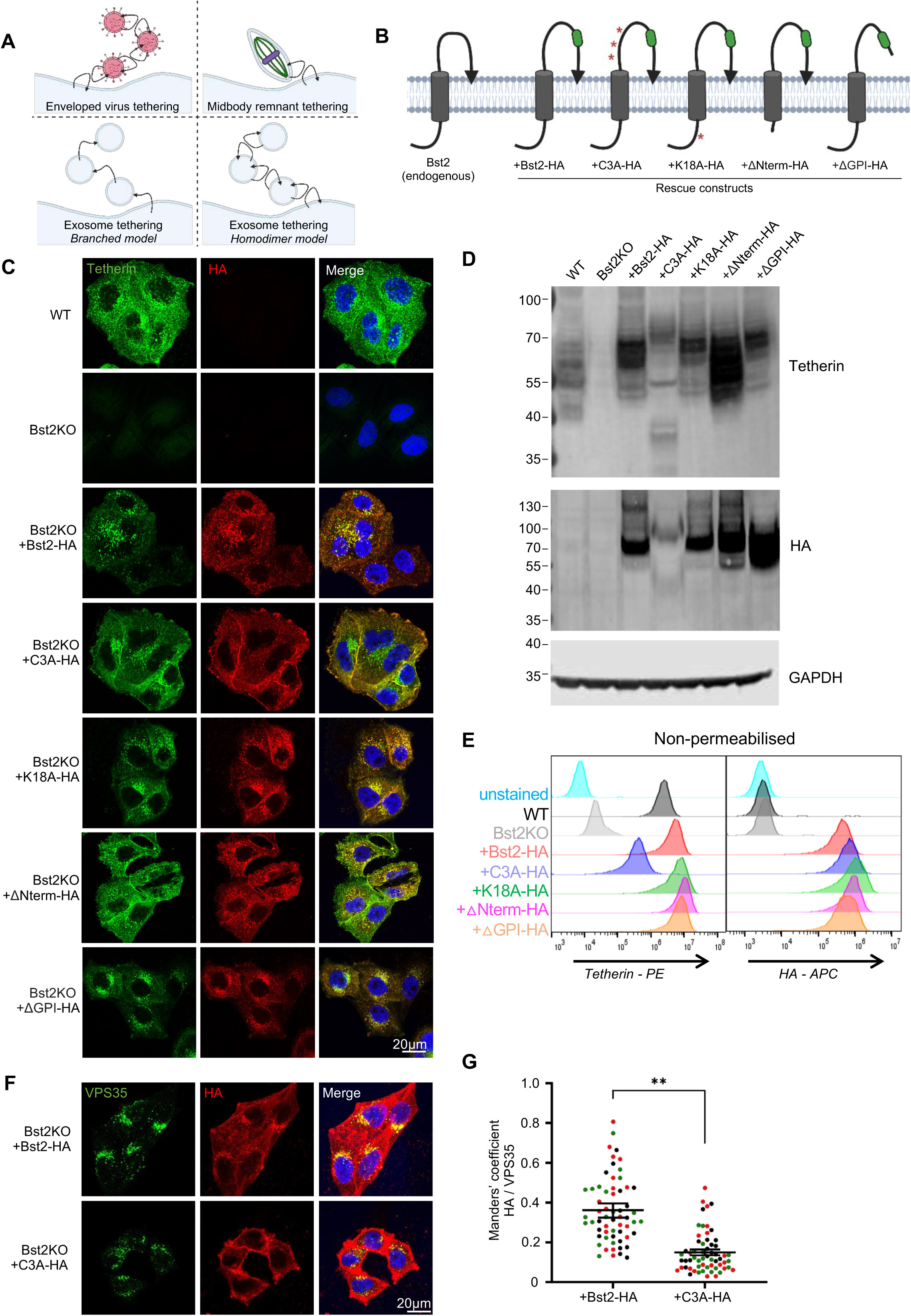
**A.** Schematic model tetherin-dependent tethering of enveloped viruses (top left), midbody remnants (top right), and potential mechanisms of exosome tethering (bottom). **B.** Diagram of tetherin membrane topology and rescue constructs. Point mutations are indicated by red asterisks. **C.** Confocal immunofluorescence images of WT, Bst2KO, and -HA rescue HeLa cells stably expressing: +Bst2-HA, +C3A-HA, +K18A-HA, +ΔNterm-HA, and +ΔGPI-HA, stained with anti-tetherin (green) and anti-HA (red) antibodies, and DAPI (blue). Scale bars = 20=μm. Representative images from three independent biological experiments. **D.** Western blot analysis of WT, Bst2KO, and -HA rescue HeLa cells stably expressing: +Bst2-HA, +C3A-HA, +K18A-HA, +ΔNterm-HA, and +ΔGPI-HA. Samples were run in non-reducing conditions (-DTT) to preserve tetherin dimerisation. Blots were probed with anti-tetherin, anti-HA and anti-GAPDH (loading control) antibodies. **E.** Flow cytometry analysis of surface tetherin expression WT, Bst2KO, and -HA rescue lines using anti-tetherin and anti-HA antibodies. **F.** Confocal images of +Bst2-HA and +C3A-HA rescue HeLa cells stained with anti-VPS35 (green), anti-HA (red), and DAPI (blue). Scale bar = 20=μm. Representative images from three independent biological experiments. **G.** Quantification of colocalisation between HA-tagged tetherin constructs and VPS35 in +Bst2-HA or +C3A-HA rescue HeLa cells using Manders’ correlation coefficients. Each dot represents an individual cell (*n* = 20 per condition); red, green, and black dots represent independent biological experiments. Bars indicate mean ± SEM (+WT-HA; 0.36 ± 0.01, C3A-HA; 0.15 ± 0.01). Statistical significance was assessed using an unpaired two-tailed *t*-test; *p* > 0.05 (ns), *p* ≤ 0.05 (*), *p* ≤ 0.01 (**), *p* ≤ 0.001(***) and *p* ≤ 0.0001 (****).

To define the molecular mechanisms of exosome tethering, we generated several tetherin mutants which we stably expressed as rescue cell lines, ectopically expressed in Bst2KO HeLa cells. Each construct was engineered with a hemagglutinin (HA) epitope inserted into the luminal domain, facilitating downstream analysis and eliminating dependence on anti-tetherin antibodies, which may not bind all mutants with equal affinity. The addition of the HA tag has been shown to minimally impact tetherin function (Billcliff et al., 2013), and Bst2-HA expression has previously been demonstrated to restore exosome tethering (Edgar et al., 2016; Palmulli et al., 2025). We established the following rescue cell lines, all of which were expressed in Bst2KO background:

**1) +Bst2-HA**: Wild-type human tetherin, modified only by the addition of the HA epitope inserted within the extracellular domain.
**2) +C3A-HA**: a mutant in which three evolutionarily conserved cysteine residues within the coiled-coil domain are replaced with alanine. These cysteine residues are essential for tetherin homodimerisation, and their mutation disrupts the ability to tether HIV-1 (Perez-Caballero et al., 2009), SARS-CoV-2 (Stewart et al., 2023), and midbody remnants (Presle et al., 2021).
**3) +K18A-HA**: Mutation of the single lysine residue in the N-terminal, short cytosolic domain. This lysine residue is targeted by Kaposi’s sarcoma-associated herpesvirus ubiquitin ligase K5, and its ubiquitination enhances tetherin trafficking to endosomes and lysosomes in an ESCRT-dependent manner (Pardieu et al., 2010).
**4)** Δ**Nterm-HA**: Deletion of the N-terminal cytosolic domain, including the YXXΦ endocytic motif and K18.
**5)** Δ**GPI-HA**: Introduction of a premature stop codon prevents the generation of the GPI anchor. The GPI promotes tetherin’s localisation to cholesterol-rich domains (Billcliff et al., 2013).

Schematics of endogenous tetherin and HA-tagged rescue constructs are shown in **Figure 2B**.

Stable expression of these constructs was confirmed by immunofluorescence microscopy (**Figure 2C**), Western blotting (**Figure 2D, Supplementary** Figure 2A), and flow cytometry (**Figure 2E, Supplementary** Figure 2B). In non-reducing conditions, tetherin appeared as a homodimer on Western blots (**Figure 2C**), confirming the loss of homodimer formation upon expression of +C3A-HA. As expected, reduction of cell lysates abolished dimer formation by cleaving disulfide bonds (**Supplementary** Figure 2A). All rescue constructs localised to some degree to the plasma membrane, to a perinuclear region and to cytosolic puncta, as shown by endogenous tetherin (**Figure 1A-D**). However, the expression of +C3A-HA displayed enhanced plasma membrane localisation (**Figure 2C, E**), consistent with previous reports (Presle et al., 2021). This redistribution may reflect the impact of tetherin monomerization on its internalisation. We hypothesise that homodimers are internalised more efficiently than monomers, as the presence of dual domains may enhance engagement with the endocytic machinery, promoting uptake of the entire complex. In contrast, monomeric tetherin likely requires each individual molecule to independently interact with endocytic adaptors, potentially reducing the overall rate of internalisation. We compared the extent of HA colocalisation with VPS35 (**Figure 2F**) – the endosomal marker that showed highest level of colocalisation with tetherin (**Figure 1A, B**) - between +Bst2-HA and +C3A-HA cells and observed a significant reduction in +C3A-HA association with endosomes relative to +Bst2-HA (**Figure 2G**).

Next, we analysed the impact of tetherin mutation on exosome tethering. Transmission electron microscopy was performed on cells to examine the capacity of each cell line to retain exosomes at their plasma membrane (**Figure 3A, Supplementary 3A**). The propensity of MVB fusion with the plasma membrane was increased by treating cells with Bafilomycin A1 (Edgar et al., 2016; Mathieu et al., 2021). Exosomes were frequently observed as clumps closely associated to the plasma membrane (**Supplementary** Figure 3A) from a number of cell lines, and clathrin-coated vesicles were frequently observed that contained multiple exosomes (**Supplementary** Figure 3B). The frequency of exosomes at the plasma membranes of cells was quantified (**Figure 3B**) and revealed that expression of +Bst2-HA, +K18A-HA, and +ΔNterm-HA restored exosome tethering to levels comparable to wild-type tetherin. In contrast, +C3A-HA expression significantly impaired exosome retention, with a frequency similar to that observed in Bst2KO cells. Cells expressing ΔGPI-HA showed a partial rescue, with more exosomes at the plasma membrane than in Bst2KO or +C3A-HA cells, but fewer than in wild-type or other functional rescue lines.

**Figure 3.**
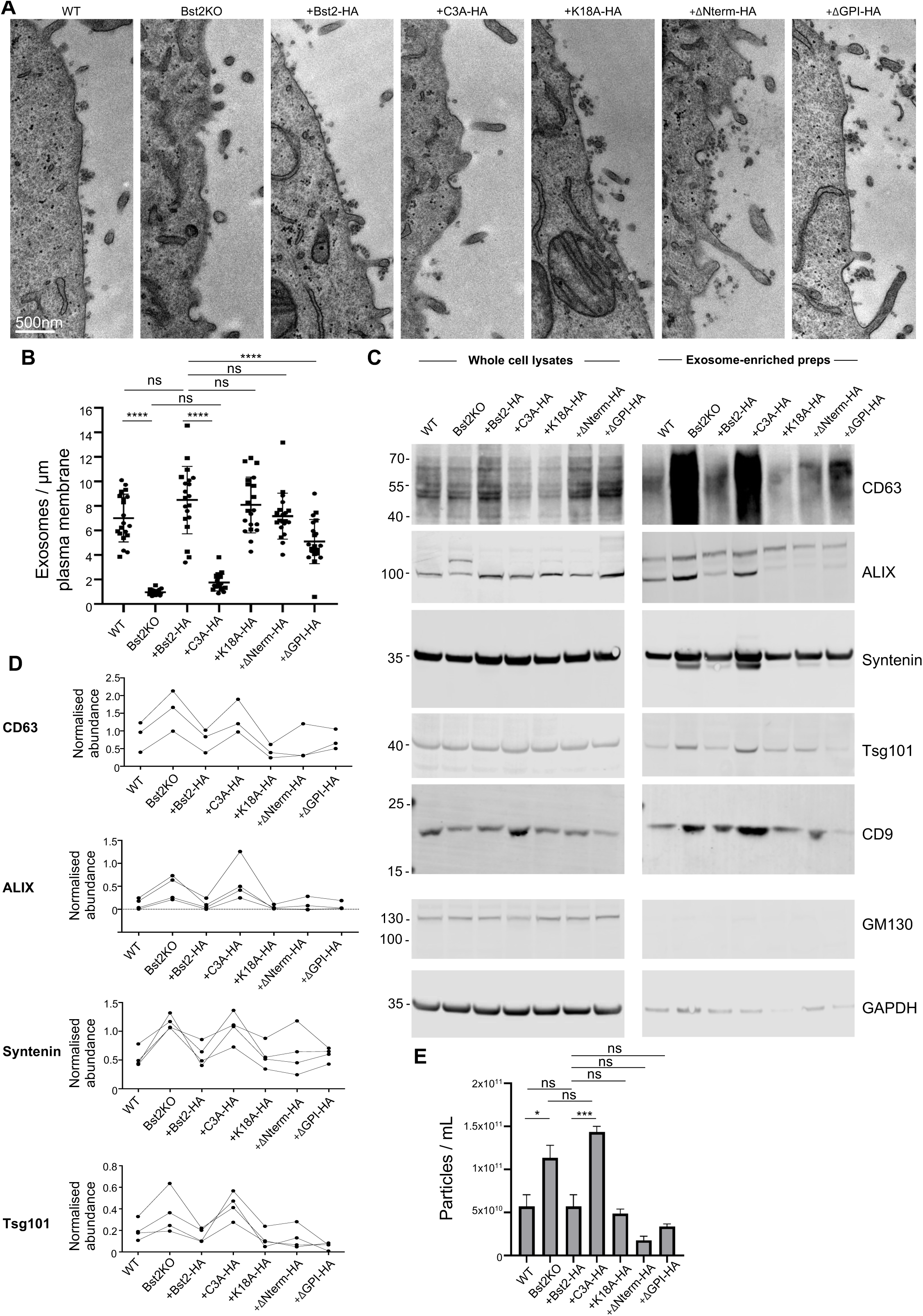
**A.** Conventional transmission electron microscopy of WT, Bst2KO, and tetherin-HA rescue HeLa cell lines that were treated with Bafilomycin A1 (100nM, 16 hr) to induce exosome release. Scale bar = 500nm. **B.** The frequency of exosomes at the plasma membrane of cells was quantified. Data shows the average number of exosomes / µm plasma membrane. Each data point represents an individual cell. *n* = two independent biological experiments, and for each independent experiment, at least two separate ultrathin sections were analysed. Bars indicate mean ± SD. Mean values: WT = 7.01 ± 1.94; KO = 0.70 ± 0.23; Bst2-HA = 8.48 ± 2.75; C3A-HA = 1.75 ± 0.68; K18A-HA = 8.09 ± 2.27; ΔNterm-HA = 7.17 ± 1.88; ΔGPI-HA = 5.10 ± 1.81. Statistical analysis was performed using one-way ANOVA with Tukey’s multiple comparisons test. *p* > 0.05 (ns), *p* ≤ 0.05 (*), *p* ≤ 0.01 (**), *p* ≤ 0.001(***) and *p* ≤ 0.0001(****). WT vs KO (*p* < 0.0001), WT vs Bst2-HA (*p* = 0.1560), KO vs C3A-HA (*p* = 0.8316), Bst2-HA vs KO (*p* < 0.0001), Bst2-HA vs C3A-HA (*p* < 0.0001), Bst2-HA vs K18A-HA (*p* = 0.9935), Bst2-HA vs ΔNterm-HA (*p* = 0.2785), Bst2-HA vs ΔGPI-HA (*p* < 0.0001). **C.** Whole cell lysates and exosome-enriched preparations were isolated from WT, Bst2KO and tetherin-HA rescue HeLa cell lines. Western blotting was performed and probed with antibodies against exosome-enriched proteins CD63, ALIX, Syntenin, Tsg101, CD9, and the EV-negative marker GM130, and loading control (GAPDH). Data representative of three independent biological experiments. **D.** Western blots were analysed and protein abundance normalised to loading controls. Relative abundance for exosome-enriched markers are shown. **E.** Quantification of particles using Nanoparticle Tracking Analysis (NTA) was performed (Zetaview). Particle concentrations are shown as particles/mL. Analysis was performed using the ZetaView system across 11 positions per sample, with ≥1000 completed tracks per condition to ensure robust quantification, and three independent biological experiments were performed. Bars indicate mean ± SEM. Statistical analysis was performed using one-way ANOVA with Tukey’s multiple comparisons test. *p* > 0.05 (ns), *p* ≤ 0.05 (*), *p* ≤ 0.01 (**), *p* ≤ 0.001(***) and *p* ≤ 0.0001(****). WT vs KO (*p* = 0.0163), WT vs Bst2-HA (*p* > 0.9999), KO vs C3A-HA (*p* = 0.377), Bst2-HA vs C3A-HA (*p* = 0.0004), Bst2-HA vs K18A-HA (*p* = 0.9959), Bst2-HA vs ΔNterm-HA (*p* = 0.1395), Bst2-HA vs ΔGPI-HA (*p* = 0.6433).

To complement this data, we examined the ability of these cells to release exosomes. Exosome-enriched preparations were isolated from conditioned media of each cell line and analysed by Western blotting (**Figure 3C, D, Supplementary** Figure 3C**, D**). The relative enrichment of the exosome-enriched proteins CD63, ALG-2 interacting protein X (ALIX), Tsg101 and syntenin, and the relative de-enrichment of cellular markers GAPDH and GM130 confirmed the successful enrichment of exosomes. Western blot analysis showed that none of the mutations impaired tetherin’s ability to traffic to exosomes, as both tetherin and -HA were present in the exosome-enriched fractions from all rescue cell lines. The levels of the exosome-enriched proteins, CD63, ALIX, Tsg101 and syntenin were elevated in preparations from Bst2KO and +C3A-HA cell lines. Quantification confirmed a statistically significant increase in exosome release between WT and Bst2KO, and also between +Bst2-HA and +C3A-HA cells (**Figure 3C, D, Supplementary** Figure 3D).

Nanoparticle analysis was performed to quantify EV particle concentrations released by each cell line (**Figure 3E**). Consistent with our biochemical data, Bst2KO and +C3A-HA cells released significantly more EVs, whereas expression of +K18A-HA, +Nterm-HA, or +GPI-HA reduced particle concentrations to levels comparable to WT or +WT-HA cells. The purity of our EV factions was analysed using DAISY analysis which allows particle-by-particle determination of both size and refractive index (Olsén et al., 2024). This makes it possible to detect and quantify subpopulations within heterogenous particle mixtures. Of particular relevance for EV studies is the identification of possible lipoprotein contaminants within samples and determination of aggregate formation, both of which can significantly alter observed particle concentrations. As a control for sample purity and consistency, we applied DAISY analysis to the EV fractions derived from our cell lines. DAISY analysis showed our samples to be consistent, with the bulk of particles having a refractive index of below 1.4 (EV reference samples have been assessed by DAISY to have refractive index <1.4 while lipoproteins have a refractive index >1.4). This increases confidence in our observed particle concentrations (**Supplementary** Figure 3E).

### Tetherin is trafficked to ILVs irrespective of ubiquitination, its GPI-anchor, or ESCRT-0 recruitment

In order for tetherin to tether exosomes, it must first be trafficked to ILVs of MVBs. We previously demonstrated that endogenous tetherin localises to the ILVs of MVBs (Edgar et al., 2016), but how tetherin is trafficked to ILVs, and whether it traffics to all subtypes of ILVs (and subsequently exosome subtypes) remains unclear.

Tetherin has two prominent features that could promote its traffic to ILVs: a cytosolic lysine residue (Lysine18), which is a target for ubiquitination (Mansouri et al., 2009), and a C-terminal GPI anchor. Both ubiquitination (Migliano and Teis, 2018) and the presence of GPI anchors (MacDonald et al., 2015) promote the traffic of cargos to ILVs.

Although a proportion of endogenous tetherin recycles from endosomes to the biosynthetic pathway (Stewart et al., 2023), the mechanism determining whether tetherin remains on the limiting endosomal membrane or is sorted into ILVs are unclear. Ubiquitination of Lysine18 drives tetherin towards ILVs, as demonstrated by the effects of the KHSV ubiquitin ligase K5 (Mansouri et al., 2009).

Cryo immunogold electron microscopy provides the ultrastructural resolution and required immunolabelling resolution to differentiate the limiting membrane of endosomes from the ILVs. Immunogold electron microscopy was performed on Bst2-HA, K18-HA and GPI-HA cell lines, and the localisation of each protein analysed using anti-HA antibodies. Consistent with prior data, Bst2-HA localised to ILVs (Edgar et al., 2016). Surprisingly, neither mutation of K18 or loss of the GPI anchor impacted tetherin localisation to ILVs (**Figure 4A**).

**Figure 4.**
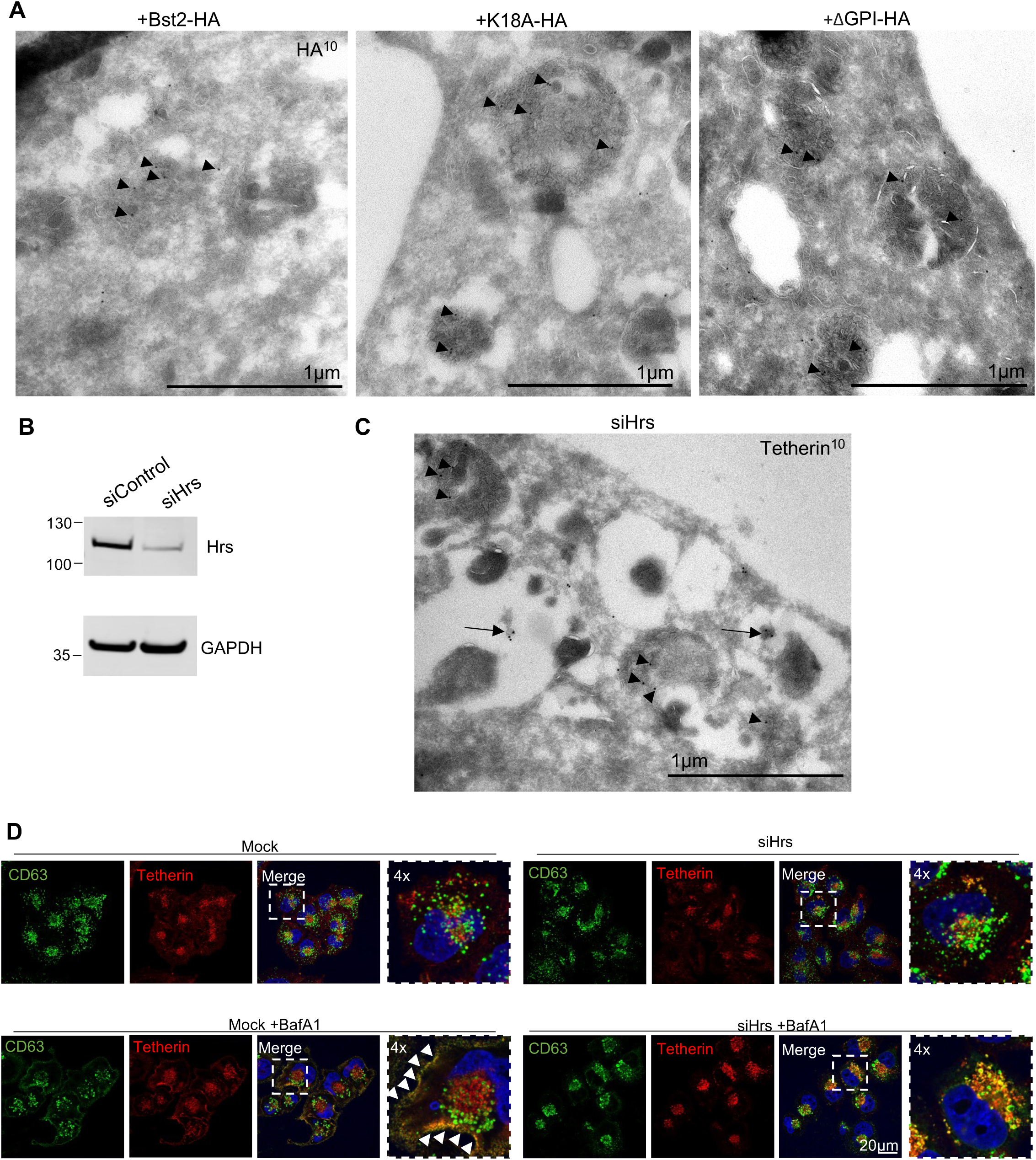
**A.** Cryoimmunogold electron microscopy was performed on + Bst2-HA, +K18A-HA, or +GPI-HA rescue cell lines. Ultrathin sections were cut, collected and stained with anti-HA antibodies and protein-A gold 10nm. Scale bar = 1μm **B.** WT HeLa cells treated with either control or Hrs-targeting siRNA. Mock and Hrs-depleted whole cell lysates were blotted for Hrs to determine its protein abundance. GAPDH is used as a loading control. Representative blots shown (from at least 3 biologically independent experiments). **C.** Cryoimmunogold electron microscopy was performed on Hrs-depleted WT HeLa cells. Ultrathin sections were cut, collected and stained with anti-tetherin antibodies and protein-A gold 10nm. Scale bar = 1μm. **D.** WT HeLa cells were depleted for Hrs using siRNA. Mock or Hrs-depleted cells were plated to coverslips and treated without or with Bafilomycin A1 (100nM, 16 hr) prior to fixation and immunofluorescence. Cells were stained using anti-CD63 (green), anti-tetherin (red) and DAPI (blue). Arrowheads highlight plasma membrane localisation of CD63 and tetherin. Scale bar = 20 μm, and 4x magnification shown.

**Figure 5.**
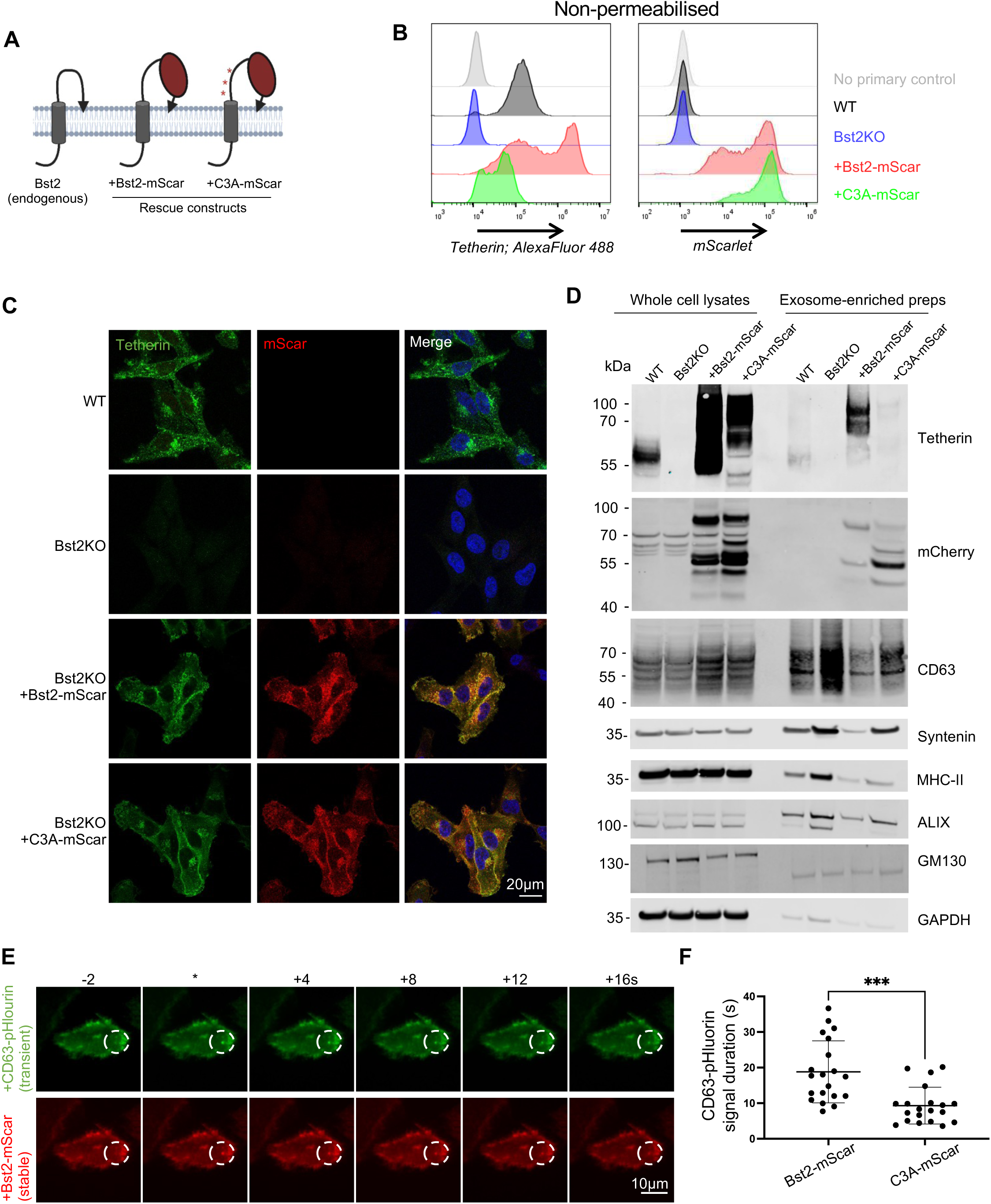
**A.** Diagram of tetherin membrane topology and -mScar rescue constructs. Point mutations within C3A-mScar are indicated by red asterisks. **B.** Flow cytometry analysis of WT, Bst2KO, Bst2KO +Bst2-mScar and +C3A-mScar MelJuSo cells. Cells were treated with interferon gamma (100 ng/mL) to promote tetherin expression. Surface tetherin (rabbit, unconjugated antibody) expression and mScarlet levels were analysed. WT cells without antibody staining was used as a no primary control. **C.** WT, Bst2KO, +Bst2-mScar and +C3A-mScar MelJuSo cells were analysed by immunofluorescence microscopy. Cells were stained with anti-tetherin (green) antibodies, and DAPI (blue). Cells were treated with interferon gamma (100 ng/mL) to promote tetherin expression. **D.** Exosome-enriched preparations were generated from WT, Bst2KO, +Bst2-mScar and +C3A-mScar MelJuSo cells. Whole cell lysates and exosome-enriched preparations were analysed by western blot and probed with anti-tetherin, anti-mCherry antibodies. The exosome enriched proteins CD63, Syntenin, MHC-II, and ALIX were analysed, as were the negative control and loading controls, GM130 and GAPDH. **E.** +Bst2-mScar MelJuSo cells were transiently transfected with CD63-pHlourin. Total internal reflection microscopy was performed. Stills from movies are shown prior to, upon (*), and following the generation of CD63-pHlourin bursts. **F.** The mean duration of CD63-pHluorin was calculated and shown. *n* = 20 total cells analysed, from 6 independent biological experiments. Bars indicate mean ± SD (Bst2-mScar: 18.99 ± 8.72; C3A-mScar: 9.33 ± 5.18). Statistical significance was assessed using an unpaired two-tailed *t*-test; *p* > 0.05 (ns), *p* ≤ 0.05 (*), *p* ≤ 0.01 (**), *p* ≤ 0.001(***) and *p* ≤ 0.0001 (****).

ILVs can be generated by a number of potential different molecular mechanisms. The ESCRT machinery is the best-described mechanism and utilises a conserved pathway of over 30 proteins to recognise and bind cargos, concentrate them to domains on the endosome limiting membrane, and subsequently drive the inward vesiculation of the limiting membrane, resulting in ILV formation. Different populations of ILVs traffic different cargos (Edgar et al., 2014; Stuffers et al., 2009; van Niel et al., 2011), and so it is likely that different exosome populations exist. It is possible that tetherin is trafficked to a subpopulation of ILVs, and as a result a subpopulation of exosomes may be tethered whilst other exosome populations may not be tethered.

The depletion of the ESCRT-0 protein, Hrs, impairs the sequestration of ESCRT-dependent cargos and limits their traffic to ILVs. Cells were depleted of Hrs (**Figure 4B**) and the localisation of endogenous tetherin analysed by immunogold electron microscopy (**Figure 4C**). Endosome enlargement and the formation of small ILVs, both characteristic of Hrs depletion (Edgar et al., 2014) were observed. Endogenous tetherin localised to ILVs in Hrs depleted cells, demonstrating that its traffic to ILVs is not solely dependent upon Hrs, and that its traffic to ILVs is not solely ESCRT-dependent.

To analyse the impact of Hrs loss on exosome tethering, we hoped to analyse exosome release biochemically. However, Hrs depleted cells displayed defects in cell division (likely due to Hrs’ requirement for cytokinesis). Instead, we analysed endosome traffic and exosome release using immunofluorescence and ultrastructural analysis. Immunofluorescence microscopy demonstrated that CD63+ endosomes remained intracellular in Hrs depleted cells and failed to relocalise to the plasma membrane upon induction with Bafilomycin A1 (**Figure 4D, Supplementary** Figure 4A). Endosomes appeared to cluster at the perinuclear region of cells in Hrs depleted cells, irrespective of treatment with Bafilomycin A1 (**Supplementary** Figure 4A**, B**). The induction of exosome release by Bafilomycin A1 has become a useful tool to study exosome release (Mathieu et al., 2021), and exosome tethering can be readily observed in mock cells treated with Bafilomycin A1 (**Supplementary** Figure 4C). However, Hrs depleted cells treated with Bafilomycin A1 did not display tethered exosomes, and instead MVBs could frequently be observed adjacent to the plasma membrane (**Supplementary** Figure 4C), indicating that Hrs-depleted MVBs were impaired in their ability to fuse with the plasma membrane.

### Live-cell imaging reveals functional tetherin impacts exosome persistence

Although we demonstrated the presence of tetherin in endosomes and ILVs, and the functional consequences of tetherin loss or mutation, we wanted to determine whether we could visualise tetherin at sites of exosome release. To achieve this, we generated mScarlet-tagged tetherin constructs and expressed these ectopically in MelJuSo cells. MelJuSo cells are widely used to study exosome release (Perrin et al., 2021), and release exosomes at steady state and without the requirement for induction via Bafilomycin A1 which would prohibit the use of CD63-pHlourin. MelJuSo cells do not express detectable levels of tetherin at steady state and so were incubated with interferon to induce tetherin expression. We stably expressed either Bst2-mScarlet or C3A-mScarlet in Bst2KO MelJuSo cells (**Figure 5A**) and verified the expression of tetherin and mScar expression by flow cytometry (**Figure 5B**) and immunofluorescence microscopy (**Figure 5C**). Stable expression of Bst2-mScar or C3A-mScar in Bst2KO HeLa cells produced mScarlet fluorescence patterns that were consistent with the localisation of both endogenous and ectopically expressed HA-tagged tetherin, as previously observed (**Figures 1A and 2C**).

Western blot analysis of whole cells confirmed the expression of both constructs, demonstrating that while Bst2-mScar retained the ability to form homodimers, C3A-mScar could not (**Figure 5D**). The functionality of mScar constructs was explored biochemically, and Bst2-mScar cells were found to be able to tether exosomes in a manner similar to WT cells, and C3A-mScar cells found to be unable to tether exosomes (**Figure 5D, Supplementary** Figure 5A**, B**) as previously demonstrated (**Figures 3A-D**).

Analogous experiments were performed in HeLa cells where Bafilomycin A1 was used to induce exosome release. The expression of mScar constructs (**Supplementary** Figure 6A**-C)** and ability to tether exosomes (**Supplementary** Figures 6D**, E**) confirmed the ability of Bst2-mScar, but not C3A-mScar, to tether exosomes. Transmission electron microscopy of HeLa cell lines also confirmed the presence of tethered exosomes from WT and Bst2-mScar cells, but with greatly reduced frequency in Bst2KO and C3A-mScar cells (**Supplementary** Figure 6F), consistent with our earlier findings using –HA-tagged tetherin (**Figure 3A-E, Supplementary** Figure 3A-D).

mScar-expressing MelJuSo cells were transiently transfected with CD63-pHluorin – a tool that allows one to visualise MVB-plasma membrane fusion (Verweij et al., 2018) – in order to allow us to examine exosome release events. Total Internal Reflection Fluorescence (TIRF) microscopy was employed to image MVB-plasma membrane fusion, and we observed the generation of CD63-pHlourin puncta alongside Bst2-mScar, confirming the delivery of Bst2-mScar and to the sites of MVB-plasma membrane fusion and exosome generation (**Figure 5E**). We examined the kinetics of CD63-pHlourin persistence between +Bst2-mScar and C3A-mScar cells, to determine whether any differences in exosome retention could be observed between exosome-competent or incompetent cells respectively by light microscopy. The duration of CD63-pHlourin signal persistence was quantified (**Figure 5F**) and revealed a significant decrease in CD63-pHlourin persistence from C3A-mScar cells when compared to Bst2-mScar cells. These findings are consistent with recent data demonstrating that functional tetherin similarly impacts exosome tethering kinetics in other cell lines (Palmulli et al., 2025).

## Discussion

Although exosomes and other extracellular vesicles can be isolated from biofluids and from conditioned media, their retention at the surface of producer cells highlights previously underappreciated functions. Whilst much research has explored the functions of released exosomes, very few studies have investigated surface-tethered exosomes. This may in-part by due to their recent discovery and the requirement for ultrastructural imaging to visualise tethered exosomes. A number of seminal exosome studies utilise transmission electron microscopy to demonstrate the presence of exosomes, and in many published images exosomes appear clustered and clumped (Carroll-Portillo et al., 2012; Colombo et al., 2014; Edgar, 2016; Johnstone, 1992; Möbius et al., 2002; Raposo and Stoorvogel, 2013; Raposo et al., 1996; Stoorvogel et al., 2002) – reminiscent of tetherin-dependent tethered exosomes.

Our data demonstrate that exosome tethering requires the formation of tetherin homodimers (**Figure 3A-D**), a requirement for the tethering of both enveloped viruses and midbody remnants. The loss of the N-terminus, or mutation of the sole cytosolic lysine residue had no impact on the ability of tetherin to traffic to endosomes (**Figure 2C**), ILVs (**Figure 4A**), exosomes (**Supplementary** Figure 3C**, D**), or their ability to tether exosomes (**Figure 3A-D**). Loss of the GPI-anchor did not impact tetherin traffic to endosomes (**Figure 2C**), traffic to ILVs (**Figure 4A**), or exosomes (**Supplementary** Figure 3C**, D**), but did functionally reduce its ability to retain exosomes (**Figure 3A-D**), consistent with the impact on virus tethering where GPI-lacking tetherin is impaired in its ability to restrict virions (Neil et al., 2008).

Exosome tethering impairs exosome release. The fate and kinetics of tethered exosomes is likely to differ between cell types relating to the relative dynamics of plasma membrane turnover. In this study, we frequently observed clusters of exosomes within clathrin-coated vesicles, consistent with their re-endocytosis upon retention at the plasma membrane (**Supplementary** Figure 3B). We also demonstrated that functional tetherin prolongs the exposure of exosomes at the cell surface (**Figure 5E, F**). Released exosomes are also likely endocytosed by neighbouring cells, although we were unable to find example of vesicles within clathrin-coated vesicles from Bst2KO or C3A-HA cells. The relative rates of endocytosis by cells, and dynamics of plasma membrane remodelling likely impact the reabsorption of exosomes. Indeed, both enveloped viruses (Mitchell et al., 2009) and midbody remnants (Presle et al., 2021) are reabsorbed by cells in a tetherin-dependent manner, demonstrating that tetherin not only prevents particle escape from cells but also promotes its reinternalisation.

Our data show that tetherin is trafficked to ILVs in the absence of cytosolic lysine or GPI-anchor (**Figure 4A**), and even in the absence of the ESCRT-0 component, Hrs (**Figure 4C**). Tetherin is trafficked through the endocytic pathway, and although it can undergo retrograde retrieval to the Golgi (Stewart et al., 2023), a proportion of tetherin remains in the endolysosomal pathway and is trafficked to late endosomes (**Figure 1A, B**). The tetherin positive late endosomes are not specifically fated for fusion with the plasma membrane with resultant formation of exosomes, as shown by the presence of tetherin in lysosomes and its accumulation upon treatment with Leupeptin (**Figure 1E-H**). Strikingly, we found that Hrs depletion impairs Bafilomycin A1-induced MVB-plasma membrane fusion and observed MVBs to be trafficked to the plasma membrane yet unable to fuse with it (**Figure 4D, Supplementary** Figure 4A-C). While Hrs depletion is known to impair cargo sorting into ESCRT-dependent ILVs - and thereby exosome biogenesis - it may also disrupt MVB-plasma membrane fusion, further impacting exosome release.

Tetherin is antagonized by enveloped viruses through distinct mechanisms, most of which limit the localisation of tetherin from the respective site of viral budding. A number of viruses promote tetherin traffic to endosomes, including HIV-1 (Mitchell et al., 2009), Kaposi’s sarcoma-associated herpesvirus (KSHV) (Mansouri et al., 2009) and SARS-CoV-2 (Stewart et al., 2023), which may promote tetherin levels on ILVs and subsequently exosomes. Viral antagonism of tetherin therefore may impact exosome fate, with implications for intracellular signalling and immune evasion.

## Funding

JRE is supported by a Sir Henry Dale Fellowship jointly funded by the Wellcome Trust and the Royal Society (216370/Z/19/Z).

## Conflict Statement

Paul Manna is an employee of Holtra AB.

## Acknowledgements

We wish to thank the electron microscopy facility at Cambridge Institute for Medical Research, University of Cambridge. The authors also gratefully acknowledge the Cambridge Microscopy Bioscience Platform for their support and assistance in this work, and thank the flow cytometry facility at the Department of Pathology, University of Cambridge.

---

**Supplementary Figure 1**

**A.** Representative confocal images showing WT HeLa cells, stained with DAPI (blue), anti-tetherin (green), and anti-LBPA (red) antibodies. Scale bar = 20=μm. **B.** Representative confocal images showing colocalisation of tetherin with TNM-BODIPY (Cholesterol) in WT HeLa cells. 4x magnification shown. Arrows highlight punctate colocalisation. Scale bar = 20=μm.

**Supplementary Figure 2**

A. Western blot of WT, Bst2KO, and -HA rescue (+Bst2-HA, +C3A-HA, +K18A-HA, +ΔNterm-HA, +ΔGPI-HA) HeLa cell lysates. Western blot was performed in reduced conditions. Blots were probed with anti-tetherin and anti-GAPDH (loading control). B. Flow cytometry analysis of protein expression in permeabilised WT, Bst2KO, and tetherin-HA rescue HeLa cell lines using anti-tetherin and anti-HA antibodies.

**Supplementary Figure 3**

A. Transmission electron micrograph of WT HeLa cells treated with Bafilomycin A1 (100nM, 16 hr). Expanded area shows plasma membrane with abundant tethered exosomes at 2x magnification. Scale bar = 10μm B. Transmission electron micrograph of WT HeLa cells treated with Bafilomycin A1 (100nM, 16 hr). Exosomes are observed within clathrin-coated pits and clathrin-coated vesicles (arrows), and expanded areas (2x magnification). Scale bar = 1μm. C. Western blot analysis of whole cell lysates and exosome-enriched preparations from WT, Bst2KO and tetherin-HA rescue HeLa cells, performed and probed with antibodies against HA and tetherin (non-reducing conditions). D. The relative abundance of CD63, ALIX, Syntenin and Tsg101 was analysed between WT and Bst2KO exosomes, and +Bst2-HA and +C3A-HA exosomes. Statistical significance was assessed using an unpaired two-tailed *t*-test; *p* > 0.05 (ns), *p* ≤ 0.05 (*), *p* ≤ 0.01 (**), *p* ≤ 0.001(***) and *p* ≤ 0.0001 (****). CD63: WT vs KO (*p* = 0.0072), Bst2-HA vs C3A-HA (*p =* 0.0263); ALIX: WT vs KO (*p* = 0.0099), Bst2-HA vs C3A-HA (*p* = 0.0298); Syntenin: WT vs KO (*p* = 0.0044), Bst2-HA vs C3A-HA (*p =* 0.0182); Tsg101: WT vs KO (*p* = 0.0376), Bst2-HA vs C3A-HA (*p =* 0.0127). E. Nanoparticle characterisation was performed on isolated EVs from each cell line using DAISY analysis. EVs were diluted in PBS and digital holographic and interferometric scattering (iSCAT) images were collected simultaneously.

**Supplementary Figure 4**

A. The particle density of CD63 was analysed between mock, mock +BafA1, siHrs, and siHrs + BafA1 treated HeLa cells. Each dot represents an individual cell (*n* = 25 cells per condition). Bars indicate mean ± SD. Mean values: mock = 9.48 ± 2.73; mock + BafA1 = 3.83 ± 1.47; siHrs = 6.61 ± 2.03; siHrs + BafA1 = 3.54 ± 1.24. Statistical analysis was performed using ordinary one-way ANOVA followed by Tukey’s multiple comparisons test. *p* > 0.05 (ns), *p* ≤ 0.05 (*), *p* ≤ 0.01 (**), *p* ≤ 0.001 (***) and *p* ≤ 0.0001 (****). Mock vs mock + BafA1 (*p* < 0.0001), mock vs siHrs (*p* < 0.0001), mock vs siHrs + BafA1 (*p* < 0.0001), mock + BafA1 vs siHrs (*p* = 0.0005), mock + BafA1 vs siHrs + BafA1 (*p* = 0.9525), siHrs vs siHrs + BafA1 (*p* < 0.0001). B. Mock or Hrs-depleted HeLa cells were treated without or with Bafilomycin A1 (100nM, 16 hours). The distribution of cholesterol (TNM-BF) and CD63 was analysed by immunofluorescence. Scale bar = 20μm. C. Transmission electron microscopy of Bafilomycin A1 treated mock or Hrs-depleted HeLa cells. The cell surface and intracellular endosome ultrastructure (arrowheads) are shown. Endosomes localised beneath the plasma membrane are shown (arrows). Images representative of two independent biological experiments. Scale bars = 1 μm.

**Supplementary Figure 5**

A. The relative abundance of CD63, syntenin or MHC-II in exosome-enriched fractions were calculated and normalised to loading controls. B. The relative protein abundance was compared between WT and Bst2KO MelJuS exosome-enriched fractions, or between +Bst2-mScar and +C3A-mScar exosome-enriched fractions. Statistical significance was assessed using a one-tailed parted *t*-test from three independent biological experiments. *p* > 0.05 (ns), *p* ≤ 0.05 (*), *p* ≤ 0.01 (**), *p* ≤ 0.001(***) and *p* ≤ 0.0001 (****).

**Supplementary Figure 6**

A. WT, Bst2KO, +Bst2-mScar, and +C3A-mScar rescue HeLa cells were analysed by confocal microscopy. Cells were fixed and stained with DAPI (blue) Scale bar = 20 μm. B. Flow cytometry analysis of surface tetherin (unconjugated rabbit antibody) and mScarlet in WT, Bst2KO, +Bst2-mScar and +C3A-mScar HeLa cells. No primary control stains represent WT cells stained only with anti-rabbit Alexa Fluor 488 secondary antibody (1:500). C. Western blotting of WT, Bst2KO, +Bst2-mScar and C3A-mScar rescue HeLa whole cell lysates. Blots were run in non-reducing conditions and probed with antibodies against tetherin, mCherry and GAPDH (loading control). D. Western blots of WT, Bst2KO, +Bst2-mScar and C3A-mScar rescue HeLa cells and exosome-enriched preparations generated from Bafilomycin A1 treated cells (100nM, 16 hr). Blots were probed with antibodies against tetherin, mCherry, and the exosome-enriched proteins CD63 (non-reducing), syntenin, ALIX, CD9, and GAPDH (loading control). E. The relative abundance of the EV-enriched proteins CD63, syntenin, ALIX and CD9 from WT, Bst2KO, +Bst2-mScar and +C3A-mScar rescue HeLa cell-derived exosome-enriched preparations were calculated relative to loading controls. F. Transmission electron microscopy was performed on WT, Bst2KO, +Bst2-mScar and +C3A-mScar HeLa cells treated with Bafilomycin A1 (100nM, 16 hr) to induce exosome release. Scale bar = 500nm.

