## Supplementary figures and images for "Exosome tethering requires tetherin homodimerisation"

### Supplemental Figure 1

#### Supplementary Figure 1

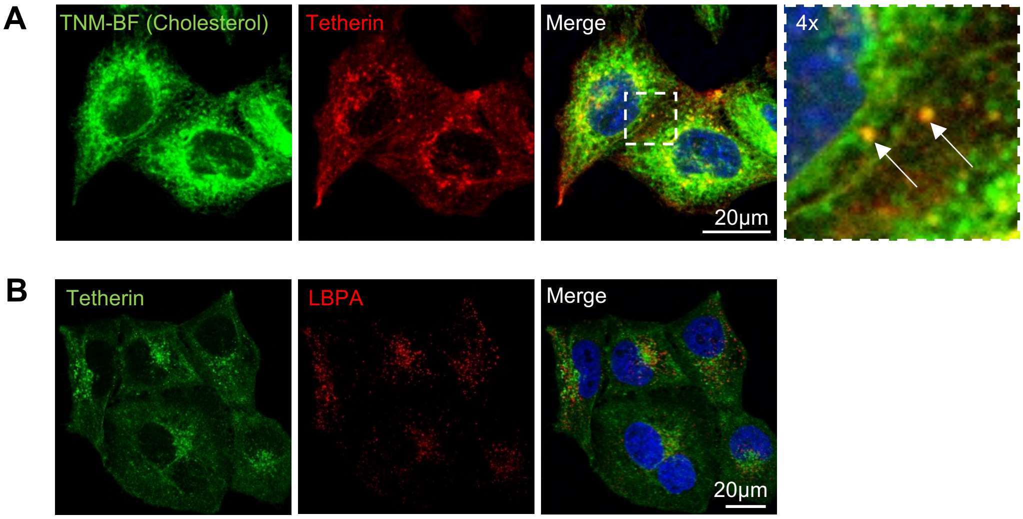

### Supplemental Figure 2

#### Supplementary Figure 2

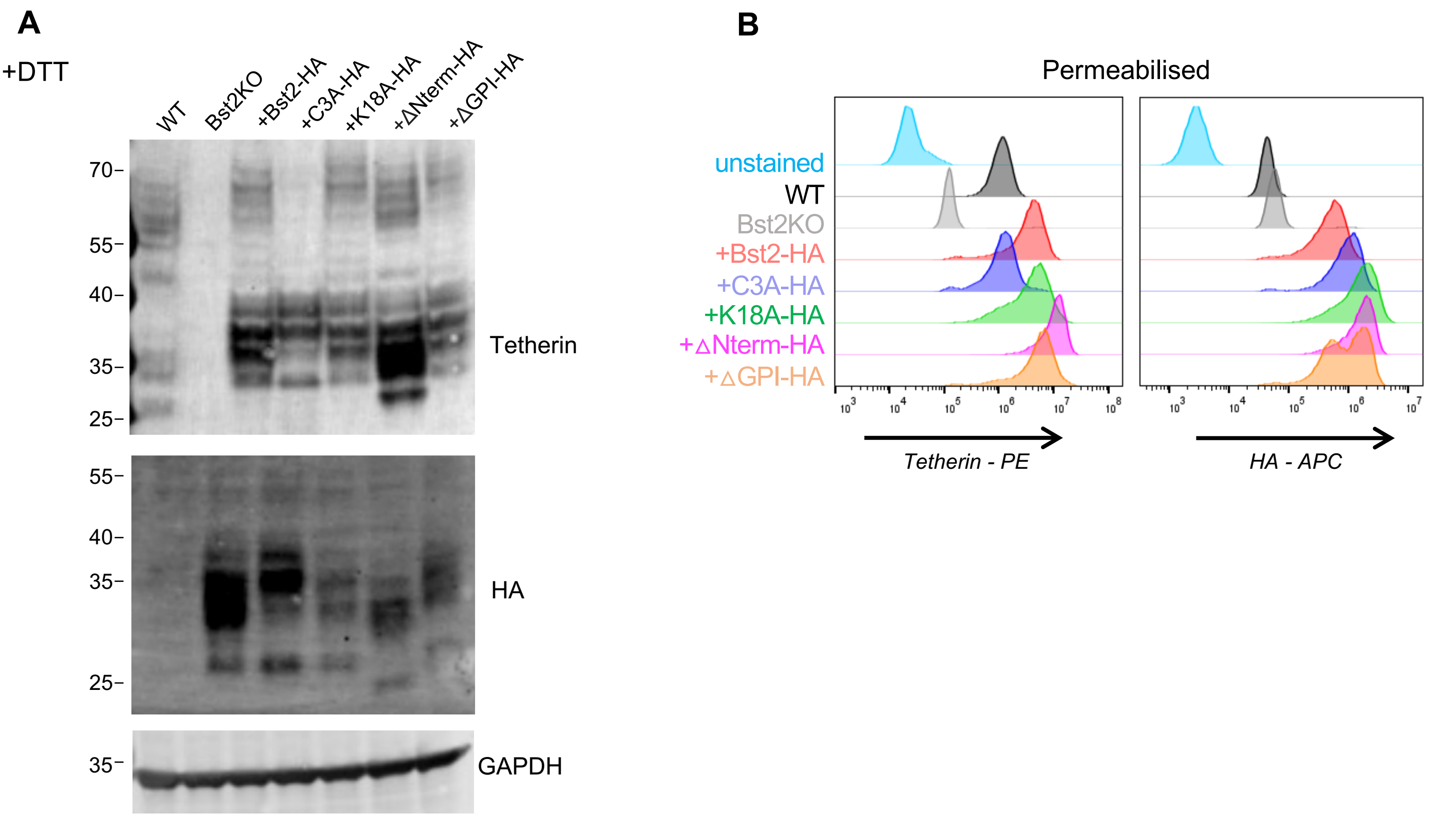

### Supplemental Figure 3

##### Supplementary Figure 3

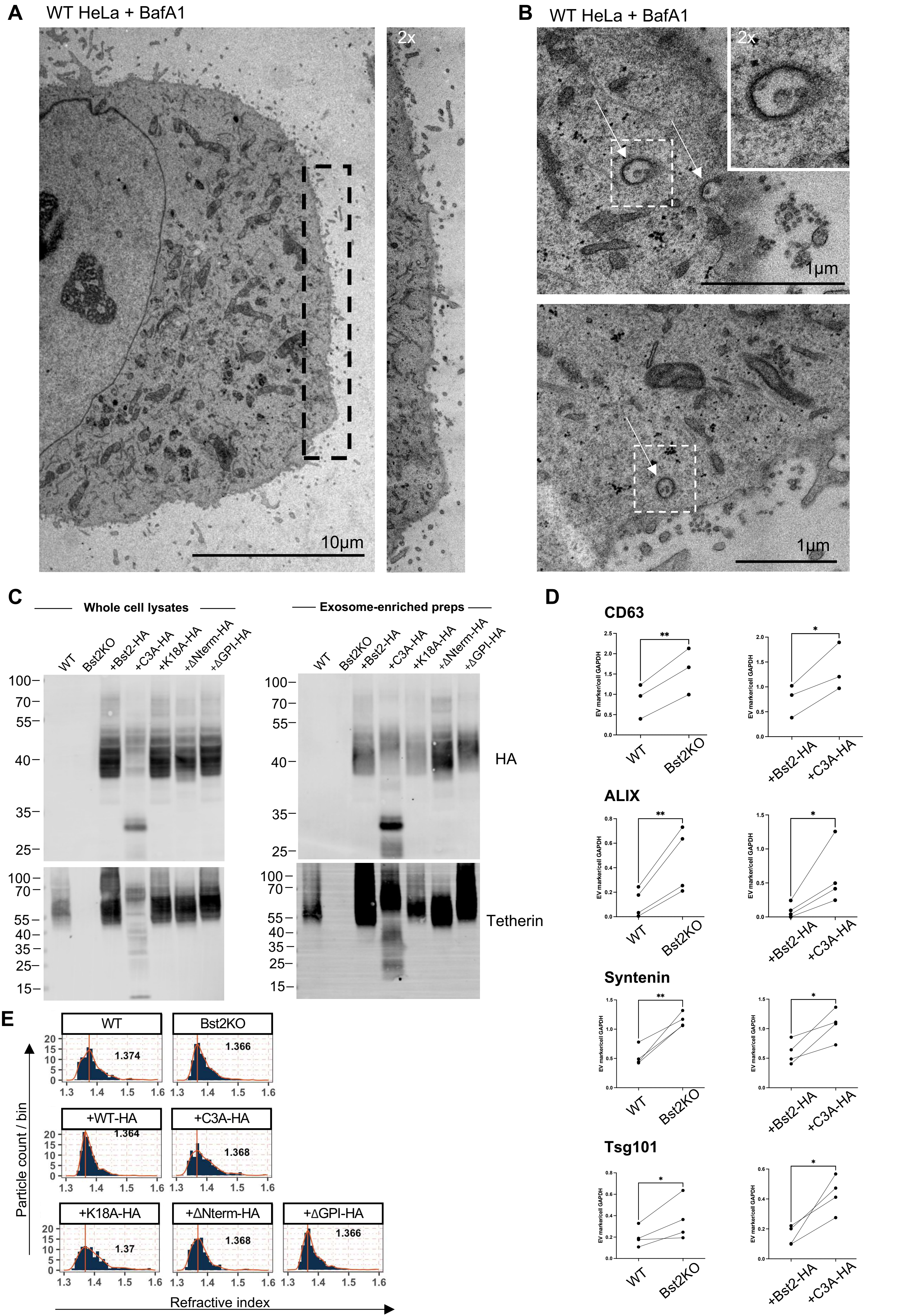

### Supplemental Figure 4

#### Supplementary Figure 4

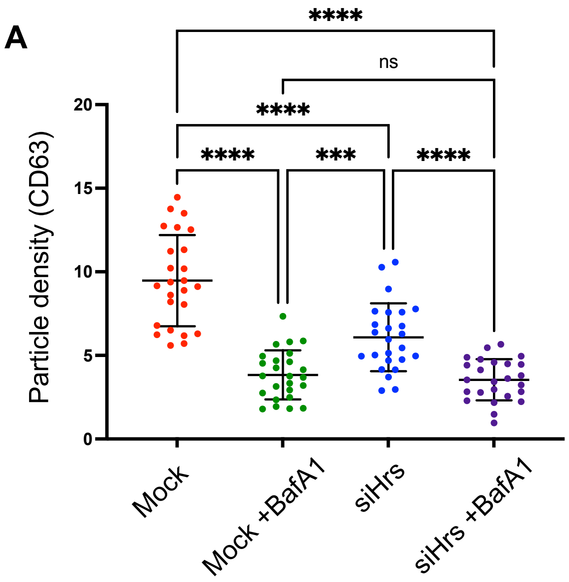


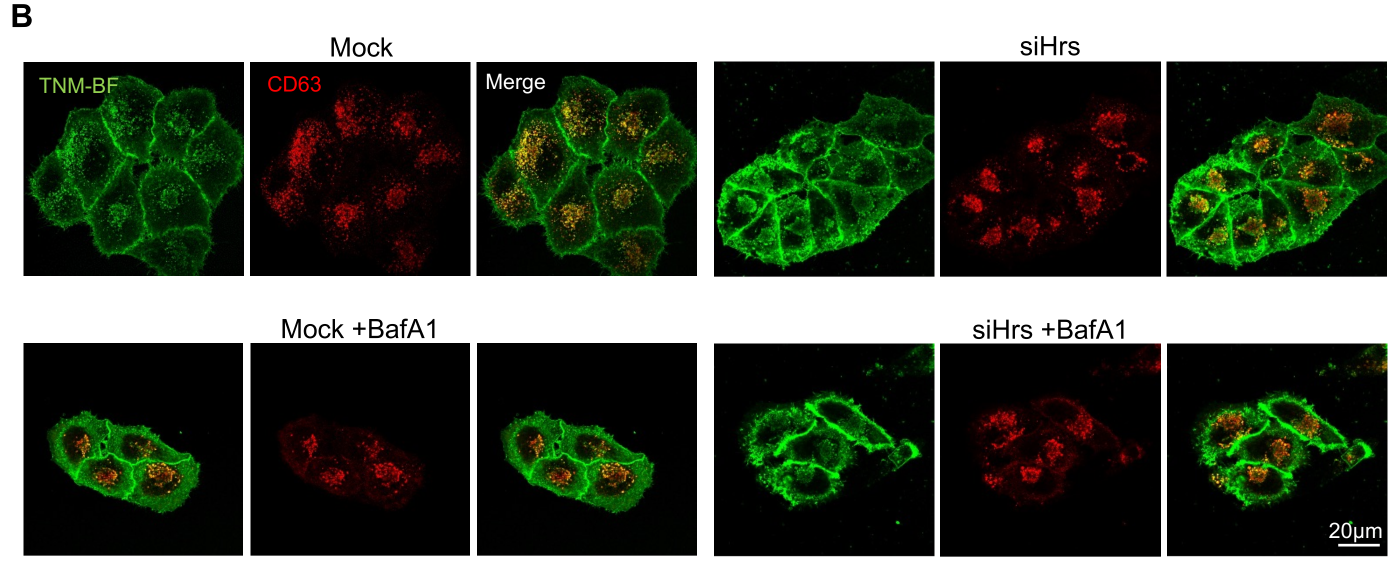


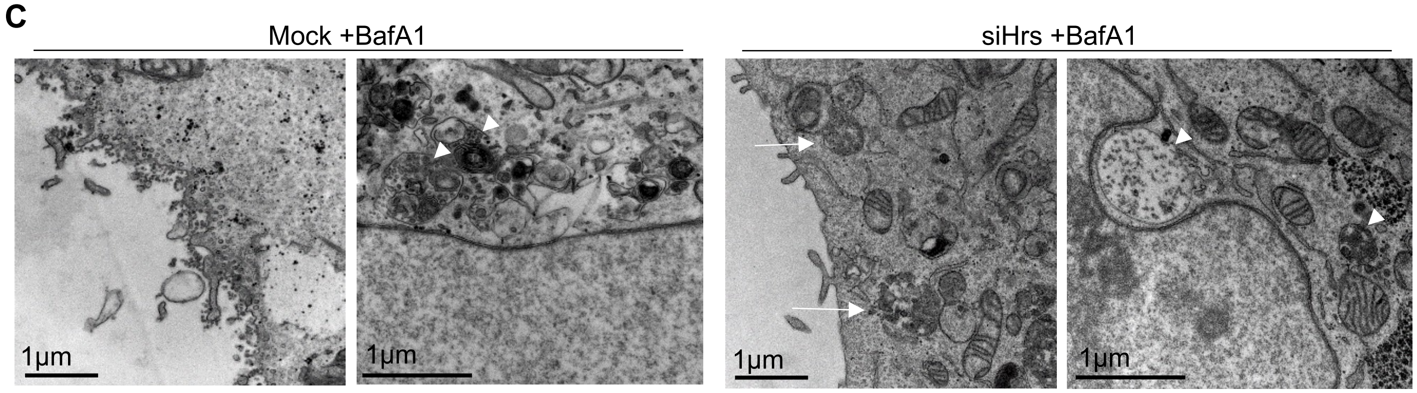

### Supplemental Figure 5

#### Supplementary Figure 5

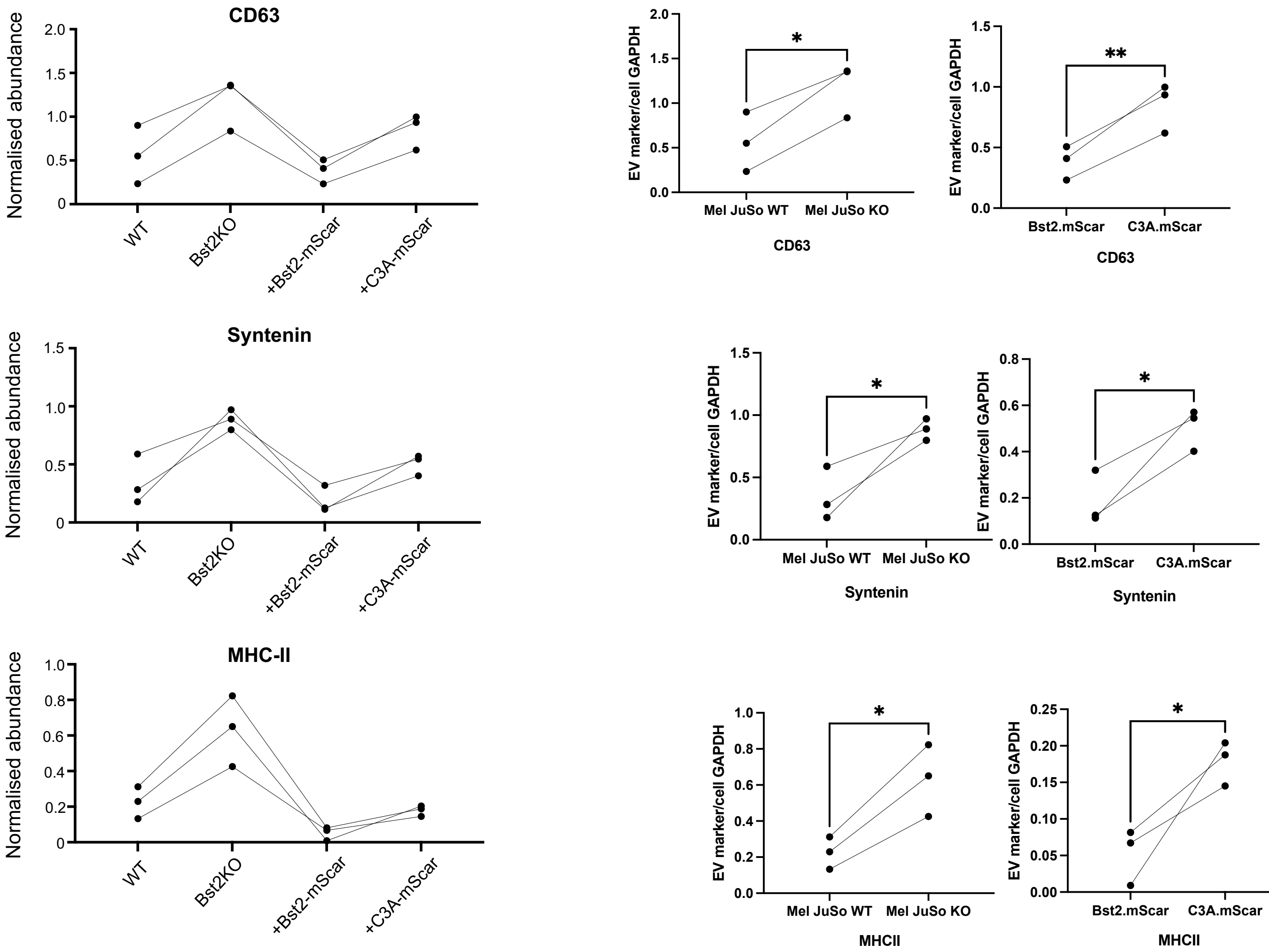

### Supplemental Figure 6

#### Supplementary Figure 6

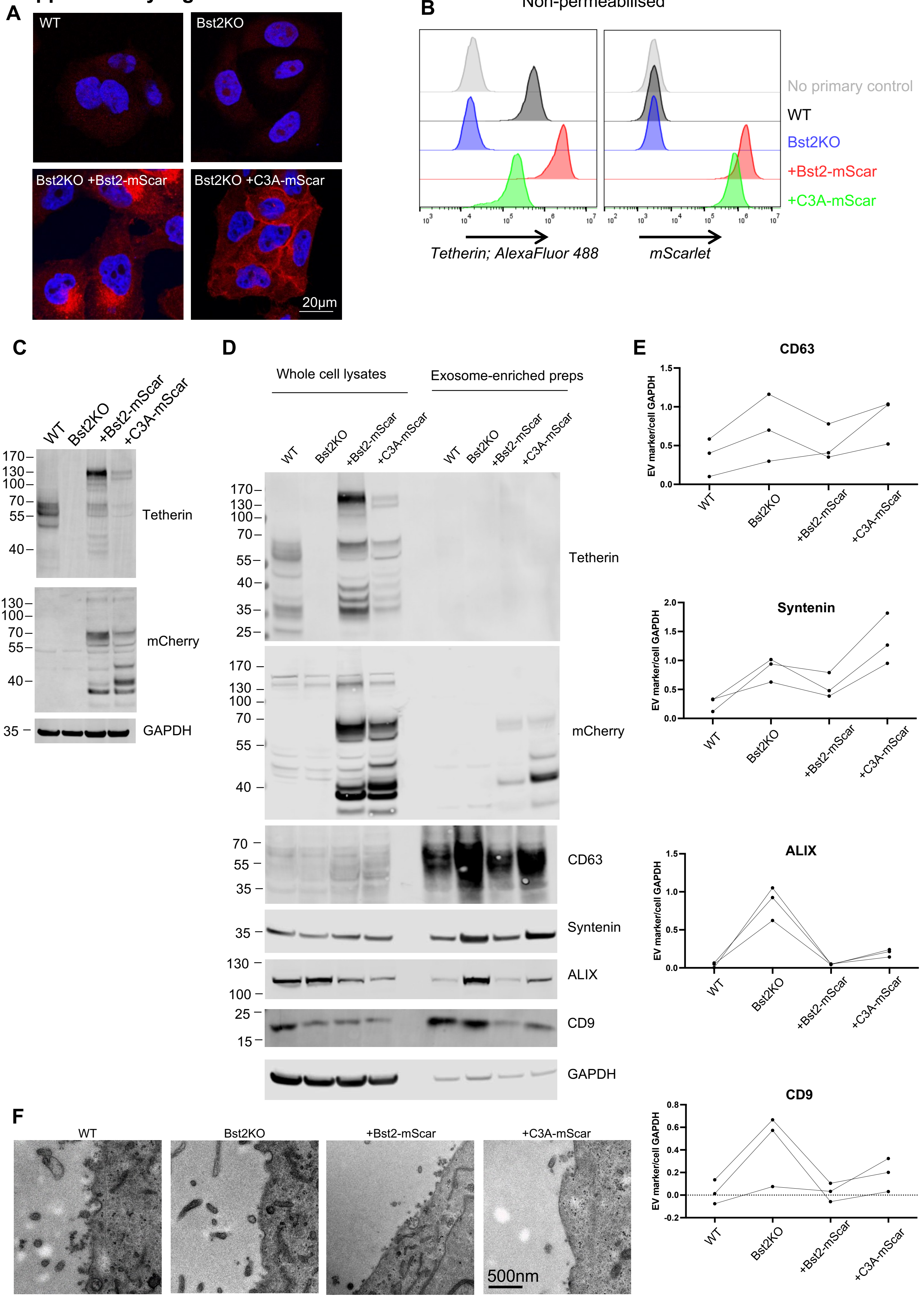
